# Branched multimeric peptides as affinity reagents for detection of α-Klotho protein

**DOI:** 10.1101/2023.01.10.523487

**Authors:** Xiyun Ye, Peiyuan Zhang, John C. K. Wang, Corey L. Smith, Silvino Sousa, Andrei Loas, Dan L. Eaton, Magdalena Preciado López, Bradley L. Pentelute

## Abstract

α-Klotho is a protein associated with aging that is expressed in the kidney, parathyroid gland, and choroid plexus. As a transmembrane protein, it acts as an essential co-receptor with the fibroblast growth factor 23 receptor complex to regulate serum phosphate and vitamin D levels. α-Klotho has an extracellular domain that can be cleaved, released and circulated in the blood stream as a soluble form. Decreased levels of α-Klotho are an indication of chronic kidney disease and other age-associated diseases. Detecting or labeling transmembrane and soluble α-Klotho is a longstanding challenge that has impeded the in-depth understanding of its role. Here we describe branched multimeric peptides that recognize α-Klotho with high affinity and selectivity in the biological milieu. The branched peptides are prepared in a single-shot synthesis by parallel automated fast-flow synthesis in under one hour. The branched α-Klotho-binding peptides show improvement in affinity relative to the monomeric versions and can be used to label Klotho for live imaging in kidney cells. Our results demonstrate the potential of automated flow technology to deliver peptide-based reagents with complex architecture and improved affinity for the selective binding of target proteins in physiological settings.

## Introduction

The aging relative protein α-Klotho (hereinafter called Klotho) is a transmembrane protein, functioning as a co-receptor of fibroblast growth factor 23 (FGF23) with fibroblast growth factor receptors 1, 2 and 4 (FGFR)^[1–4]^. The FGFR/Klotho/FGF23 complex regulates phosphate and calcium homeostasis at the kidney. Klotho has an extracellular domain composed of two tandem domains^[2]^. Cleavage of the extracellular domains produces a soluble form of Klotho which is released into the circulation and is found in the blood, urine, and cerebrospinal fluid^[4]^. Decreased membrane-bound or soluble Klotho protein levels are correlated with aging-related dysfunction, including diabetes, chronic kidney disease, cancers, and cardiovascular diseases^[4]^. Design of reagents with high affinity and selectivity to detect Klotho can help understand both Klotho distribution in the body to facilitate early detection of aging-associating diseases and guide the development of anti-aging therapies.

Disruption of protein-protein interactions (PPIs) are challenging targets for molecular recognition as it usually involves multivalent, and cooperative interactions between two proteins to form tightly bound complexes^[5]^. Peptides can mimic the native binding epitope at PPI regions, thus capturing features of the extended molecular interaction^[6–8]^. Peptide-based affinity reagents are endowed with the advantages of low molecular weight (<10 kDa), suitability for large-scale manufacture by solid-phase peptide synthesis, and enhanced stability in complex biological media. With these advantages, peptide reagents can be improved upon to penetrate cell membranes and have potential longer half-life and superior circulation profiles *in vivo*^[9–11]^.

The interaction interface between Klotho and FGF23 provides opportunities for designing PPI-disrupting peptides targeting Klotho. The FGF23 N-terminus interacts primarily with FGFR, while its C-terminus engages Klotho ^[2,12,13]^ (Figure 1A). The DiMarchi group identified peptides that mimic the C-terminus of FGF23 based on the minimal binding domain determined from the Klotho crystal structure^[14]^ (Figure 1B). These peptides were able to antagonize activity of FGF23 in cells. We selected a top performing peptide from this published work (**Pep-10**: PMASDPLGVVRPRARM) and measured the binding affinity between **Pep-10-Biotin** (Figure S1) and recombinant mouse Klotho (mKlotho) using bio-layer interferometry (BLI). **Pep-10-Biotin** was immobilized onto the streptavidin tips, and serial dilutions of mKlotho were incubated with the tips to measure binding and dissociating events, giving an apparent dissociation constant (KD) of 2.3 ± 0.5 μM (Figure. S2A). The binding affinity of **Pep-10** is insufficient to detect Klotho in serum which is at nanomolar range. However, **Pep-10**serves as a promising starting point for development of peptide-based affinity reagents.

**Figure 1.**
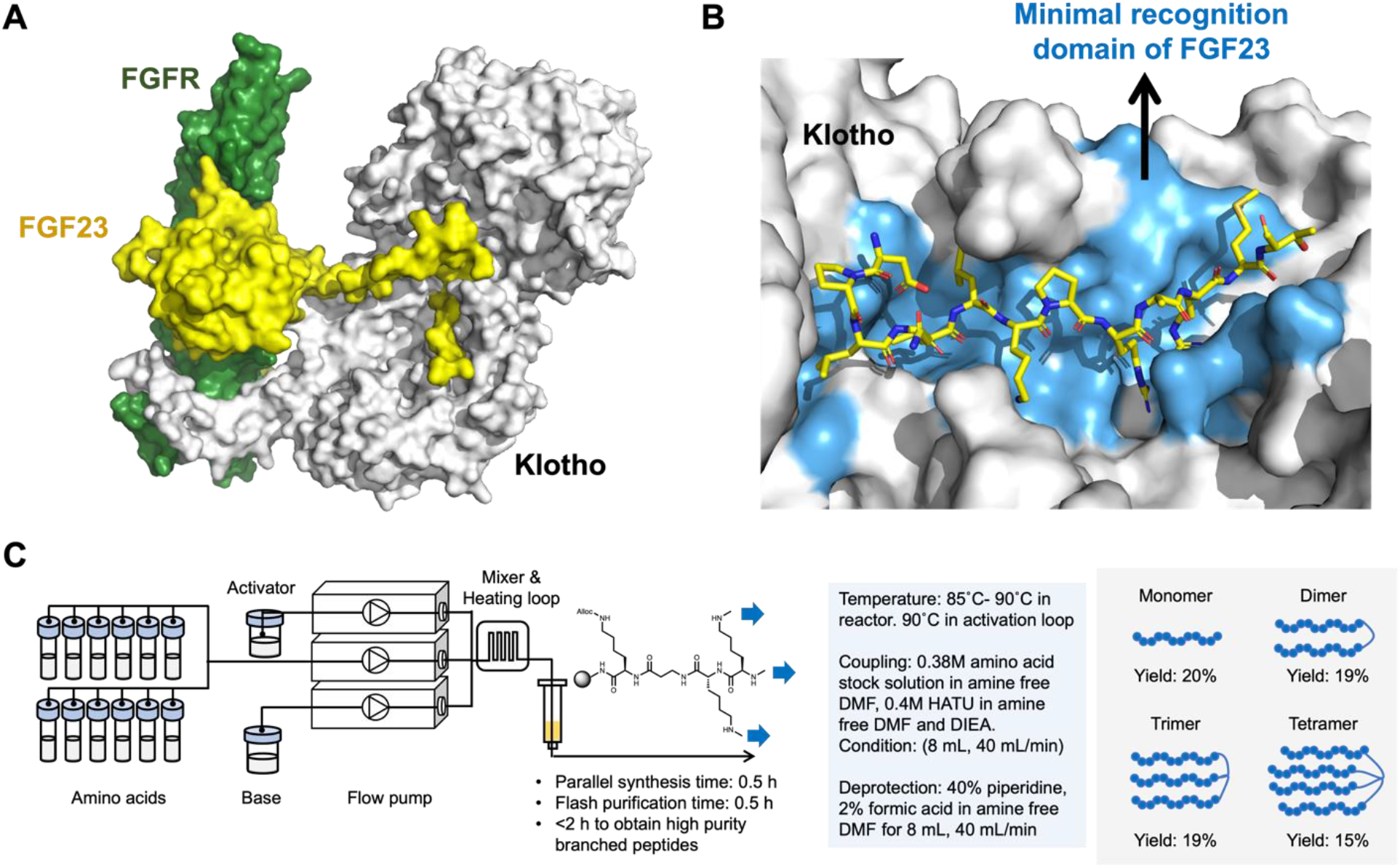
Automated flow synthesis provided access to high-affinity peptide reagents for the detection of Klotho. (A) Structure of the protein-protein interaction between co-receptor protein Klotho, receptor protein FGFR and signaling protein FGF23, modified from PDB:5W21. Klotho is in white, FGFR protein is in green and FGF23 is in yellow. (B) Visualization of the binding pocket of Klotho and the C-terminus of FGF23. FGF23^188-200^ is shown as sticks and the Klotho van der Waals protein surface in white. The contacting surface of Klotho is colored in blue. (C) Schematic representation of an automated flow protein synthesizer (AFPS) coupled to a flash peptide chromatography system to parallelly synthesize branched peptides.

## Results and Discussion

Preparation of multimeric, branched forms of peptide reagents has been used to maintain selectivity while increasing avidity, which allows for stronger binding compared to the correspondent monomeric peptides^[15–17]^. However, the synthesis of branched peptides can be challenging due to the shared branching amino acid. Employing fast-flow synthesis with our in-house high-temperature automated synthesizer, we were able to synthesize Klotho-binding branched peptides within half an hour, ranging from dimeric to tetrameric variants^[18–20]^ (Figure 1C). During the flow synthesis, each step fluorenylmethyloxycarbonyl (Fmoc) deprotection is monitored by in-line UV-Vis spectroscopy to ensure each synthesis cycle proceeds with high efficiency. Notably, the C-terminal residue of **Pep-10** is Met which introduces steric hindrance when elongating multiple peptide chains. To reduce the possibility of Met oxidation and potentially increase the synthesis yield, we used **KB1** (PMASDPLGVVRPRARA) instead of **Pep-10** (PMASDPLGVVRPRARM) as monomer for branched peptides. **KB1-Biotin** has similar K_D_ as **Pep-10-Biotin** in the BLI assay as 2.6 ± 0.5 μM (Figure S2B). We accomplished synthesis of each branch of the multimeric Klotho-binding peptides simultaneously through a parallel elongation process. After global deprotection and cleavage, the crude material was purified by high performance flash chromatography (HPFC) and freeze-dried to obtain the pure branched peptides. For each synthesis batch, we obtained ~4 μmol of pure material (16 mg of dimer **KB2**, 24 mg of trimer **KB3**, and 32 mg of tetramer **KB4**) in ~20% yield starting from 20 μmol of Rink amide resin (Figure 1C).

**Figure 2.**
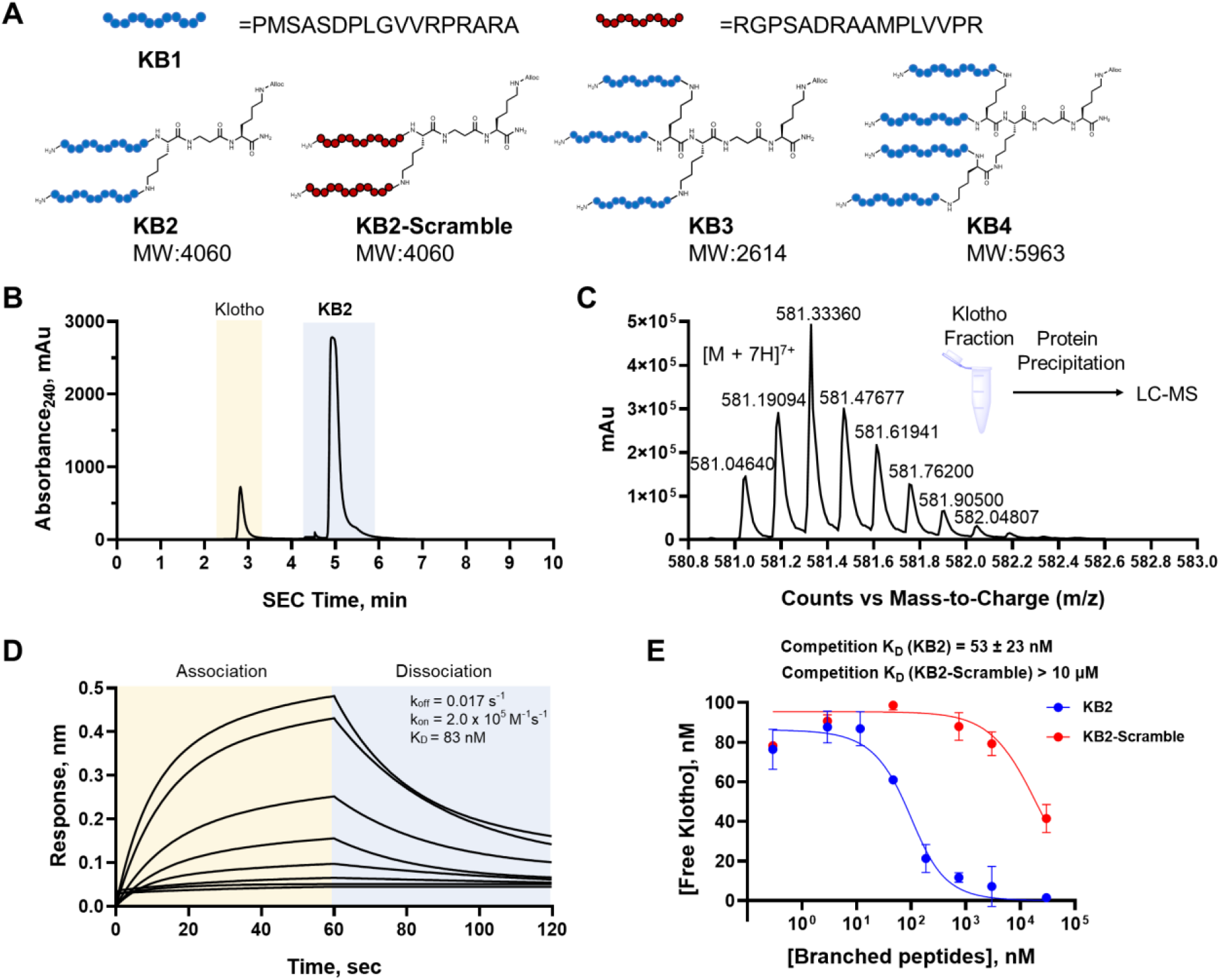
Biophysical studies characterized the molecular interactions between branched multimeric peptide binders and Klotho. (A) Sequence of monomeric Klotho-binding peptide (**KB1**), dimeric Klotho-binding peptide (**KB2**) and the control scramble peptide (**KB2-Scramble**), trimeric Klotho-binding peptide (**KB3**) and tetrameric Klotho-binding peptide (**KB4**). (B) HPSEC-MS analysis of the Klotho–**KB2** complex. HPSEC achieved a base-line separation of the protein (1 μM, retention time 2.7 min) and peptide (10 μM, retention time 4.8 min) fractions. (C) The mass spectrum of **KB2** was detected in the protein fractions (peak with retention time of 2.7 min in **Figure 2B**) after precipitation of Klotho, indicating a strong peptide-protein interaction. (D) Analysis of the Klotho–**KB2-Biotin** interaction through a direct binding measurement by BLI. **KB2** was functionalized with Biotin-PEG4 using the preinstalled Alloc-handle to provide **KB2-Biotin**, which was further immobilized onto streptavidin tips. The solutions contained various concentrations of Klotho as 1000, 500, 250, 125, 62.5, 31.3, 15.6, 7.8 nM. The direct binding was recorded by a BLI system to study the dissociation and association kinetics (n = 2). The off-rate of **KB2-Biotin** was estimated to be 0.017 s^-1^ and the on-rate was estimated to be 2.0 × 10^5^ M^−1^ s^−1^. (E) A competition assay by BLI was established to estimate the competitive apparent dissociation constant (K_D_) in a label-free format (n = 2). Unlabeled **KB2** binds to Klotho with K_D_ = 53 ± 23 nM. The control peptide **KB2-Scramble** does not significantly associate with Klotho (K_D_ > 10 μM). Error bars indicate SD.

To investigate whether branched multimeric peptides display improved binding affinity to Klotho, we applied analysis by high-performance size exclusion chromatography-mass spectrometry (HPSEC-MS), direct binding measurements by BLI, competitive binding assays by BLI, and fluorescence polarization (FP) assays. These biophysical assays demonstrated 100-fold enhanced binding affinity towards Klotho by branched multimeric peptides compared to monomer **KB1**.

HPSEC-MS analysis directly assesses the formation of the peptide-protein complex. Unlabeled peptides were individually mixed with Klotho and partitioned by HPSEC to separate protein-bound and unbound fractions. The eluted fraction corresponding to Klotho was subsequently analyzed by mass spectrometry (MS) to confirm the recovery of bound branched peptides. **KB2** (recovery yield 4%, defined as the amount of peptide detected in the eluted Klotho fraction divided by total amount of peptide before mixed with Klotho), **KB3** (recovery yield 3%), and **KB4** (recovery yield 2%) were retained in the Klotho fraction in three replicates (Figures 2B, 2C, and S3). Notably, the parent **KB1** monomeric peptide was not observed in the Klotho fraction, indicating a potentially weaker interaction leading to dissociation of the complex on the HPSEC column. HPSEC-MS qualitatively showed enhanced binding to Klotho by the branched peptides relative to monomeric **KB1**.

Direct binding measurements by BLI were completed using **KB2-Biotin**, **KB3-Biotin** and **KB4-Biotin**, in a similar manner to **KB1-Biotin**. These biotinylated probes were immobilized and immersed into solutions containing various concentrations of mKlotho as 1000, 500, 250, 125, 62.5, 31.3, 15.6, 7.8 nM. The kinetic parameters, k_on_ (association rate constant) and k_off_ (dissociation rate constant), were then determined based on curve fitting and the K_D_ was calculated as k_off_/k_on_. Compared to the monomer **KB1** (K_D_ = 2.6 ± 0.5 μM), the branched peptides **KB2**, **KB3** and **KB4** gave K_D_s of 83 ± 42 nM, 35 ± 15 nM and 88 ± 56 nM, which resulted in a 27.7, 65.7 and 26.1-fold improvement of binding affinity respectively (Figure 2D and S2).

Competitive binding assays against the parent **KB1** binder allowed us to determine competition K_D_ values by BLI and confirmed Klotho site-specific binding by the branched multimeric variants. **KB1-Biotin** was immobilized onto the streptavidin biosensor tips. Increasing concentration of unlabeled **KB1** through **KB4** were mixed with a constant concentration of mKlotho (100 nM) to measure binding and dissociating events. **KB1-Biotin**-immobilized tips were immersed into each of these solutions to estimate the fraction of unbound mKlotho and interpolate the competition K_D_ values. **KB1** competed off binding of mKlotho by **KB1-Biotin** and gave a competition K_D_ as 6.5 ± 1.2 μM (Figure S4A). **KB2** (K_D_ = 53 ± 23 nM), **KB3** (K_D_ = 75 ± 29 nM) and **KB4** (K_D_ = 139 ± 43 nM) displayed progressively lower affinity (higher competition K_D_ values) with the increasing number of branches (Figures 2E and S4). Within the three branched peptides, the highest improvement in binding affinity was observed with the dimer **KB2** in the competitive binding assays. To assess the binding selectivity of **KB2**, we synthesized a control dimeric branched peptide based on a scrambled **KB2** sequence, named **KB2-Scramble. KB2-Scramble** returned a high competition K_D_ (> 10 μM) in the competitive binding assay (Figure 2E), indicating weak binding to mKlotho with at least 200-fold higher competition KD. Therefore, **KB2-Scramble** was used as a negative control of **KB2** in downstream assays. Two additional proteins, MDM2 and 12CA5 were also used to evaluate binding selectivity of **KB2**. Biotinylated probe **KB2-Biotin** displayed no binding to these two proteins in the direct binding measurements by BLI (Figure S5). These results indicated the branched peptides showed non-promiscuous and site-specific binding interactions with Klotho.

We labeled **KB2** with the fluorophore TAMRA (5-carboxytetramethylrhodamine) and determined its apparent dissociation constant K_D_ to be 154 ± 24 nM, estimated by an FP assay (Figure S6). This orthogonal assay suggested fluorophore-labeled **KB2** retains nanomolar binding affinity to Klotho. The binding affinities of **KB2-TAMRA** and **KB2-Scramble-TAMRA** were also evaluated in the competitive binding assays by BLI to provide a comparison to the unlabeled molecules (Figure S4D). The competition K_D_ of **KB2-TAMRA** was determined as 57 ± 26 nM, which is similar to **KB2** (K_D_ = 53 ± 23 nM). As in the case of **KB2-Scramble** alone, **KB2-Scramble-TAMRA** displayed minimal binding to Klotho (K_D_ > 10 μM). In summary, these findings ensured binding between **KB2** and Klotho is not affected upon modification with TAMRA.

To transition from the mKlotho form used in the above binding assays to human Klotho (hKlotho), we measured the direct binding of **KB1-Biotin** to hKlotho by BLI. The experimentally determined K_D_ was 3.0 ± 0.6 μM, similar to the value determined for **KB1-Biotin** binding to mKlotho (K_D_ = 2.6 ± 0.5 μM) (Figure S7A). Furthermore, **KB2** bound hKlotho with K_D_ = 85 ± 22 nM in the competitive binding assay by BLI (Figure S7B). This value is comparable to the apparent dissociation constant determined against mKlotho (K_D_ = 53 ± 23 nM) in the same assay (Figure 2E). Similarly, **KB2-TAMRA** displayed K_D_ = 90 ± 31 nM to hKlotho while **KB2-Scramble-TAMRA** returned a K_D_ > 10 μM, indicating the attachment of TAMRA also has no effect on **KB2**’s binding to hKlotho (Figure S7C and S7D).

Taken together, these biophysical assays suggested the lead branched dimer peptide **KB2** has an enhanced binding affinity towards Klotho compared to monomeric **KB1**, for example in the competition assays: competition K_D_ of **KB1** is 6.5 μM which is 123-fold higher than competition K_D_ of **KB2** that is 53 nM. Our flow synthesis technology delivered peptides with diverse branching architectures including dimeric peptide **KB2** had the highest Klotho binding affinity, and therefore was selected for downstream target engagement assays.

To study engagement of Klotho in a complex cellular environment by **KB2**, we first investigated whether KB2 can capture supplemented recombinant mKlotho from cell lysates. We added recombinant mKlotho into the cell lysate of CaSki cells, a human cervical cancer cell line that was reported to display minimal Klotho expression^[21]^ (Figure S8), to a final concentration of 100 nM. Klotho was significantly enriched in the pulled-down fraction by **KB2-Biotin** starting at 200 nM, consistent with its binding affinity determined by BLI (Figure 3A). The **KB1-Biotin** and control probe **KB2-Scramble-Biotin** did not enrich mKlotho even at 1 μM (Figure 3A and S9). To confirm the protein enriched from lysate was Klotho, we ran a control pull-down with a commercially available Klotho antibody (KM2076). The pull-down fraction by **KB2-Biotin** and by Klotho antibody gave bands with the same molecular weight, confirming it was Klotho being enriched (Figure S9). In a self-competition assay, pull-down of Klotho by **KB2-Biotin** could be competed off by unlabeled **KB2** added to the cell lysate in a dose-dependent manner, but not by **KB2-Scramble** (Figure 3B). Together, the dose-dependent and the self-competition pull-down assays show direct mKlotho engagement by **KB2** in cell lysate. To investigate selectivity of pull-down by **KB2-Biotin**, silver stain was performed on the pull-down eluents. mKlotho was the only significantly enriched protein by **KB2-Biotin** at 200 nM but not control **KB2-Scramble-Biotin**, demonstrating selective pull-down achieved by **KB2-Biotin** (Figure S10). Next, we evaluated hKlotho engagement in human kidney 2 (HK-2), a proximal tubular cell line that expresses detectable levels of Klotho endogenously^[22]^. **KB2-Biotin** enriched hKlotho from HK-2 cell lysate at 200 nM, but not **KB2-Scramble-Biotin** (Figure 3C). These studies demonstrate direct binding of **KB2** to both recombinant mKlotho and endogenous total hKlotho, supporting the notion that the branched multimeric peptide has the potential to selectively engage its target in biological milieu.

**Figure 3.**
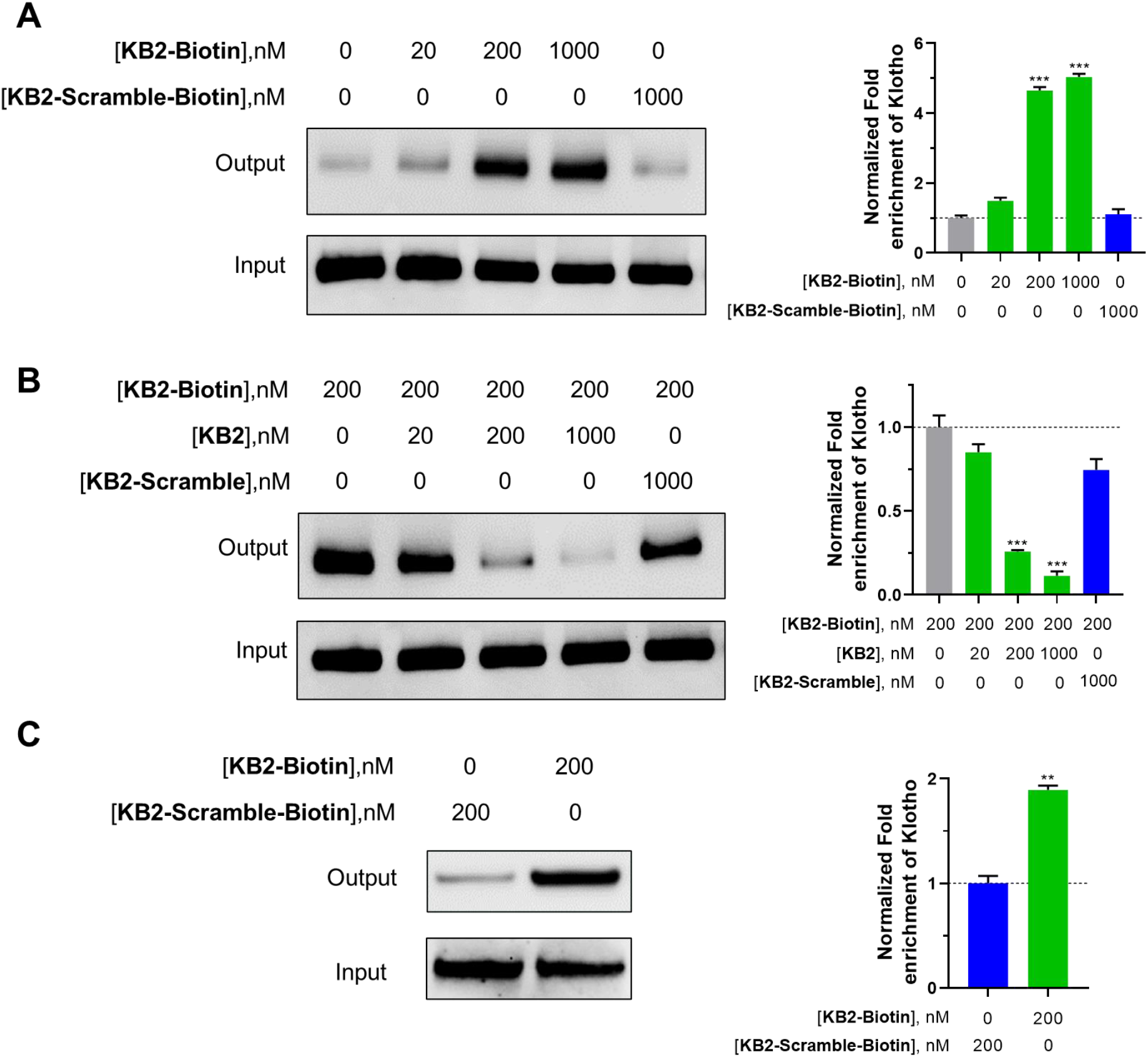
Pull-down assays from human cell lysates showed direct Klotho engagement by KB2. (A) Pull-down probe **KB2-Biotin** enriched recombinant mKlotho from 100 μg of CaSki cell lysate in a dose dependent manner. Control probe **KB2-Scramble-Biotin** displayed no enrichment of Klotho at 1 μM. The bar graph depicts quantification of protein pull-down (n = 2). (B) Pull-down of mKlotho by **KB2-Biotin** can be competed off by **KB2** in a dose-dependent manner, but not by **KB2-Scramble**. The bar graph depicts quantification of protein in the competitive pull-down assay (n = 2). (C) **KB2-Biotin** significantly enriched endogenous hKlotho from HK-2 cell lysate at 200 nM. The bar graph depicts quantification of protein pull-down (n = 2). **P < 0.01, ***P < 0.001, as determined by ANOVA. Error bars indicate SD.

Labeling specific proteins of interest in live cells remains a challenge due to the general dearth of stable, selective affinity reagents that are not antibody-based. Since **KB2-TAMRA** selectively binds to Klotho with high affinity and selectivity, proved by the competition assay by BLI (Figure S4D), we used it as our lead Klotho affinity reagent for fluorescence microscopy imaging of HK-2 cells. A time course study was performed to optimize the imaging protocol, suggesting a strong intracellular fluorescence signal was obtained in fixed HK-2 cells upon incubation with 500 nM of **KB2-TAMRA** for only 5 min (Figure S11). The scrambled control, **KB2-Scramble-TAMRA** did not display appreciable fluorescence emission in HK-2 cells, indicating minimal binding to Klotho and potential low internalization efficiency into cells (Figure S11). In these assays, the cell membrane was demarked with the use of the membrane-specific dye Wheat Germ Agglutinin (WGA) Alexa Fluor™ 350 Conjugate. To validate that intracellular fluorescence comes from the labeling of Klotho by **KB2-TAMRA**, we performed several target engagement and competition studies using recombinant hKlotho and unlabeled **KB2** binder (Figure 4). Pre-incubation of **KB2-TAMRA** with rhKlotho significantly decreased the intensity of the fluorescence signal, consistent with a reduced amount of free, unbound **KB2-TAMRA** available for cell labeling (Figure 4). We also validated that **KB2-TAMRA** and **KB2** engage the same binding site in cells by a competition experiment. Pre-treatment of cells with **KB2** off-competed labeling of endogenous Klotho by **KB2-TAMRA**, resulting in significantly decreased intracellular fluorescence (Figure 4). This indicates that **KB2-TAMRA** and **KB2** bind the same site of Klotho in cells, consistent with the *in vitro* binding assay (Figure 2E and S7C).

**Figure 4.**
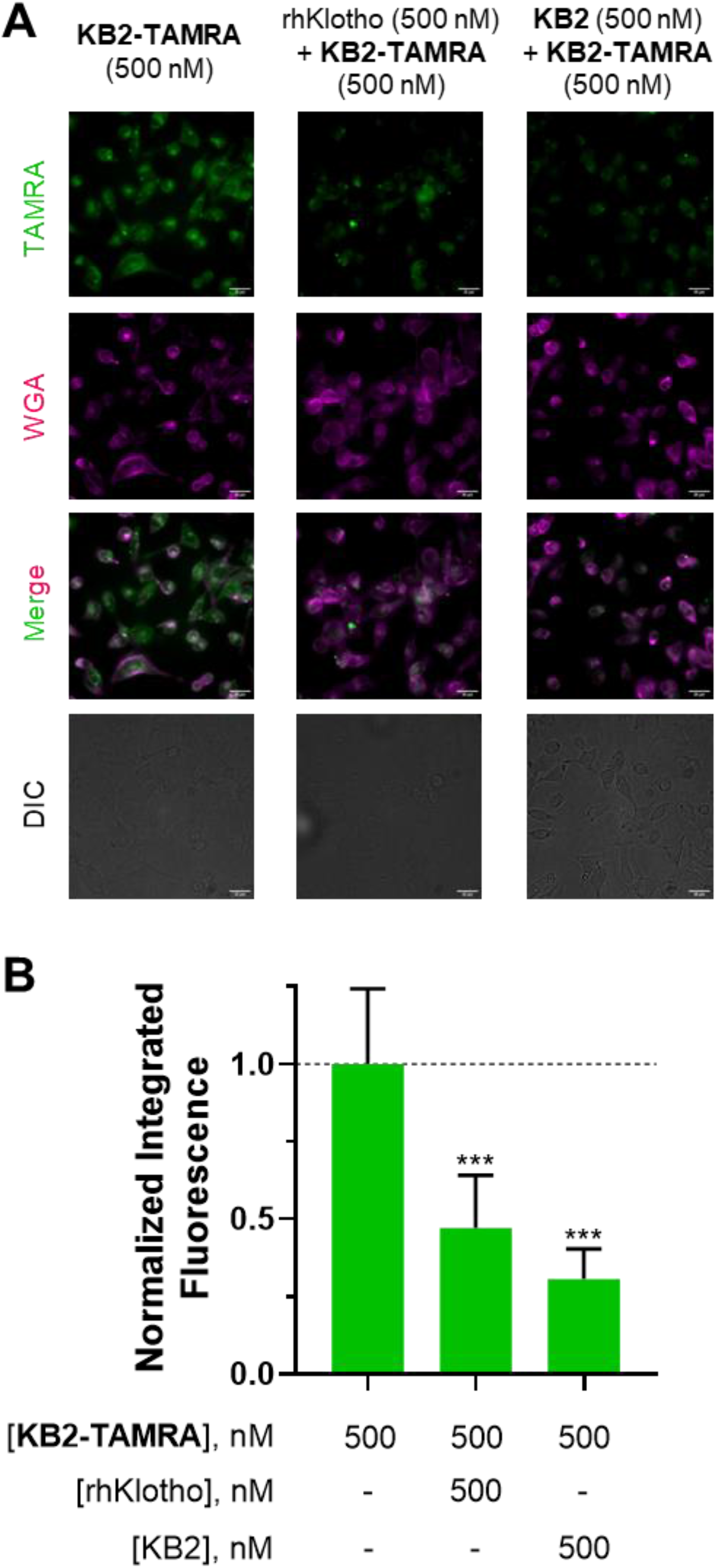
Fluorescence microscopy imaging revealed target engagement of KB2-TAMRA labeling total Klotho in fixed HK-2 cells. (A) Left: detection of intracellular fluorescence and potential labeling of endogenous total hKlotho following 5 min of co-treatment with **KB2-TAMRA** at 500 nM and Wheat Germ Agglutinin (WGA) Alexa Fluor^™^ 350 Conjugate at 25 ng/μL. Fixation of cells was performed after the treatment. Fluorescence of **KB2-TAMRA** was acquired in the orange channel (λ_ex/em_ = 542/597 nm). Fluorescence of WGA Alexa Fluor^™^ 350 was acquired in the blue channel (λ_ex/em_ = 390/435 nm). Middle: HK-2 cells treated for 5 min with a mixture of pre-incubated **KB2-TAMRA** (500 nM) and recombinant hKlotho (500 nM) displayed significantly decreased intracellular fluorescence. Right: HK-2 cells were pre-treated with **KB2** (500 nM) for 1 h followed by co-treatement of **KB2-TAMRA** at 500 nM and WGA at 25 ng/μL. The pre-treatment of KB2 also displayed significantly decreased intracellular fluorescence. (B) The bar graph depicts quantification of normalized integrated fluorescence in panel A (n = 10). ***P < 0.001, as determined by ANOVA. Error bars indicate SD. Scale bar = 25 μm.

To further validate the selective binding, we knocked down endogenous Klotho levels in HK-2 cells with a pool of siRNAs (KL siRNA, Figure S12), followed by treatment with **KB2-TAMRA**. Mock transfection had no effect on fluorescence in cells (Figure 5A) as expected. Notably, knocking down Klotho levels by KL siRNA reduced the fluorescence of **KB2-TAMRA**, supporting that fluorescent labeling by **KB2-TAMRA** is indeed Klotho-dependent (Figure 5).

**Figure 5.**
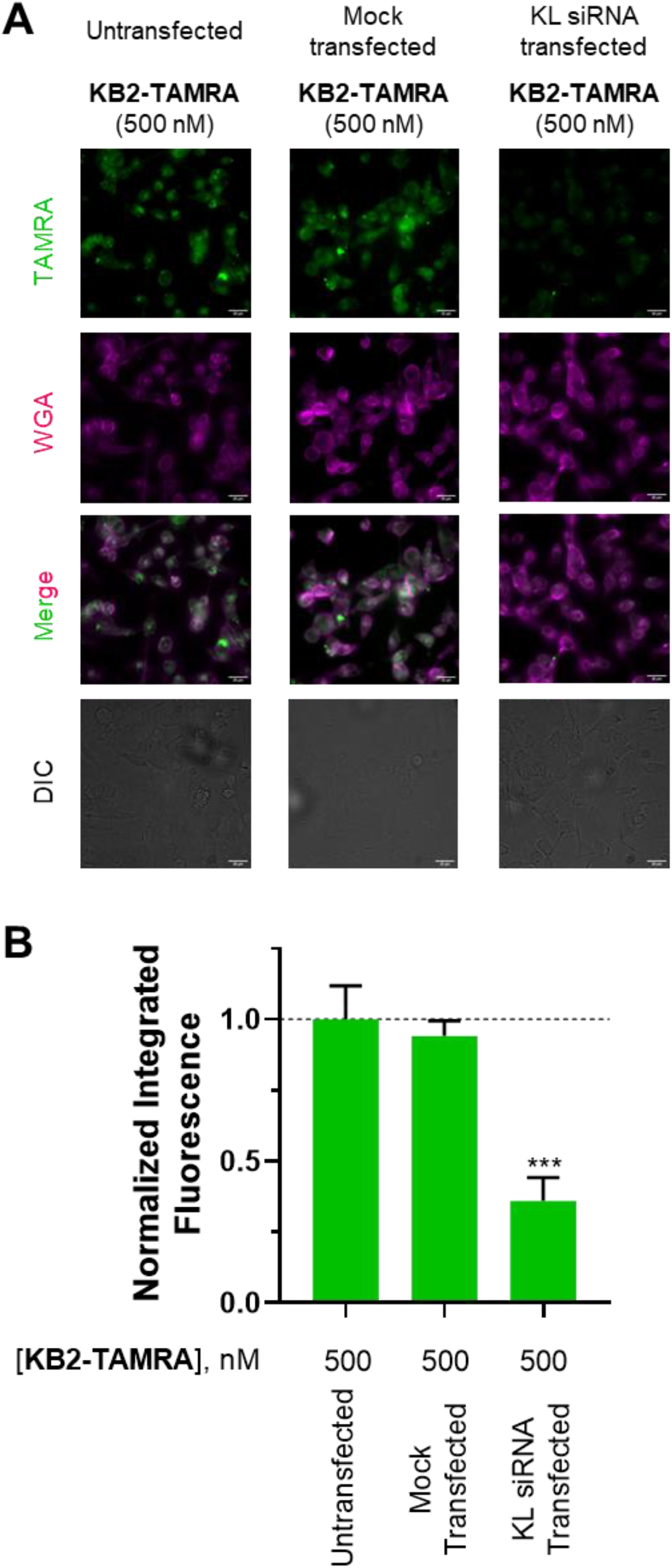
Validation of mode of action of Klotho labeling by KB2-TAMRA in fixed HK-2 cells. (A) In the first 2 columns, HK-2 cells were untransfected or mock transfected by RNAiMAX reagents. In the last column, HK-2 cells were transfected with KL siRNA by RNAiMAX reagents. After transfection, **KB2-TAMRA** at 500 nM was co-treated with Wheat Germ Agglutinin (WGA) Alexa Fluor^™^ 350 Conjugate at 25 ng/μL in HK-2 cells. Fixation of cells was performed after the treatment. Fluorescence of **KB2-TAMRA** was acquired in the orange channel (λ_ex/em_ = 542/597 nm). Fluorescence of WGA Alexa Fluor^™^ 350 was acquired in the blue channel (λ_ex/em_ = 390/435 nm). Scale bar = 25 μm. (B) The bar graph depicts quantification of normalized integrated fluorescence (n = 5). ***P < 0.001, as determined by ANOVA. Error bars indicate SD. Scale bar = 25 μm.

We also investigated whether **KB2-TAMRA** can internalize and engage its target in a different cell line with lower Klotho expression levels. We incubated **KB2-TAMRA** with Klotho-deficient CaSki cells (Figure S8) using a similar protocol to HK-2 cells. Under these conditions, **KB2-TAMRA** exhibited a significantly lower fluorescence intensity in CaSki cells relative to Klotho-expressing HK-2 cells, potentially indicating reduced cell penetration and consistent with the expected lower Klotho levels (Figure S13A).

**KB2-TAMRA** selectively labels Klotho in Klotho endogenously expressed cells. We compared the fluorescence microscopy images of **KB2-TAMRA** with Klotho antibody (Figure S13B) under the same conditions and found the fluorescence patterns to be similar, suggesting selective labeling of Klotho by **KB2-TAMRA**. To monitor Klotho subcellular localization, we used DAPI to stain the cell nuclei (Figure S14). Pearson correlation coefficients were used to quantify the co-localization of nucleus dye DAPI versus **KB2-TAMRA** (Figure S14A) and Klotho antibody (Figure S14B). Low correlations (The coefficient between DAPI and **KB2-TAMRA** is 0.29. The coefficient between DAPI and Klotho antibody is 0.35) indicated **KB2-TAMRA** and the Klotho antibody stained the membrane as well as cytosol and intracellular organelles of HK-2 cells, but not the nucleus. These results support membrane and cytoplasmic localization of Klotho consistent with the literature^[23]^. The Klotho antibody requires overnight incubation, followed by additional permeation steps and incubation with the secondary antibody, which makes it unsuitable for live cell labeling. On the contrary, our fluorescent peptide **KB2-TAMRA** was able to label Klotho in live HK-2 cells due to a short incubation time (5 min) and mild conditions (Figure S15). These imaging experiments support that **KB2-TAMRA** can be used as an efficient and selective probe to potentially label Klotho in both fixed and live cells.

## Conclusion

In brief, we report a flow synthesis strategy to deliver branched peptides as a high-affinity material and selective probe to engage Klotho in a cellular context. Their binding affinities were validated by multiple biophysical and biochemical assays. These branched peptides can be developed into fluorescent labeling reagents with other potential uses. To further optimize the Klotho-targeting peptides, we envision a rational design strategy to affinity-mature the peptides, leading to improved biophysical properties. Optimization to further improve the affinity, reduce molecular weight, and improve stability in vivo may pave the way for discovery of next-generation Klotho labeling agents.

## Corresponding Author

**Bradley L. Pentelute** – Department of Chemistry and Center for Environmental Health Sciences, Massachusetts Institute of Technology, Cambridge, Massachusetts 02139, United States; The Koch Institute for Integrative Cancer Research, Massachusetts Institute of Technology, Cambridge, Massachusetts 02142, United States; Broad Institute of MIT and Harvard, Cambridge, Massachusetts 02142, United States; orcid.org/0000-0002-7242-801X; Email

**Magdalena Preciado López** – Calico Life Sciences LLC, San Francisco, CA 94080, United States of America, Email

## Acknowledgement

Financial support for this work was provided by Calico Life Sciences (to B.L.P.). We acknowledge support from the Swanson Biotechnology Center Microscopy Core Facility at the Koch Institute for Integrative Cancer Research at MIT through the use of their epifluorescence microscope (NCI Cancer Center Support Grant P30-CA14051).

## Conflict of interest

B.L.P. is a co-founder and/or member of the scientific advisory board of several companies focusing on the development of protein and peptide therapeutics. A provisional patent disclosure was filed regarding the methodology and compounds described in this study.

## Entry for the Table of Contents

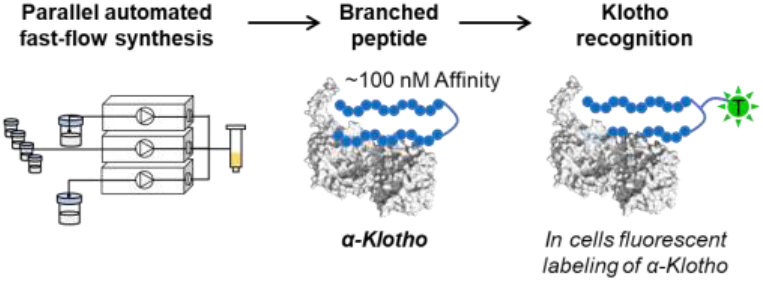
α-Klotho recognition. Branched peptides that recognize α-Klotho are prepared via parallel automated fast-flow synthesis. The branched peptide binders show at least 30-fold improvement in affinity relative to the monomeric versions and can be used to label Klotho for live imaging in kidney cells.

