## Supporting information for "Branched multimeric peptides as affinity reagents for detection of α-Klotho protein"

[2] Calico Life Sciences
1170 Veterans Blvd., South San Francisco, CA 94080 (USA)

[3] AbbVie Bioresearch Center
100 Research Dr, Worcester, MA 01605 (USA)

[4] The Koch Institute for Integrative Cancer Research, Massachusetts Institute of Technology
500 MainStreet, Cambridge, MA 02142 (USA)

[5] Center for Environmental Health Sciences, Massachusetts Institute of Technology
77 Massachusetts Avenue, Cambridge, MA 02139 (USA)

[6] Broad Institute of MIT and Harvard
415 Main Street, Cambridge, MA 02142 (USA)

[#] These authors contributed equally to this work.
*To whom correspondence should be addressed:



**Contents**

1. Supplementary figures ... 3
2. Methods and Materials ... 19
   2.1 Materials ... 19
   2.2 Mouse Klotho (mKlotho) protein production ... 19
   2.3 Fast flow synthesis of branched peptides ... 19
   2.4 Manual preparation of peptidyl resins with linker 1,2, and 3 ... 19
   2.5 Automated flow peptide synthesis (AFPS) set-up ... 20
   2.6 Linker coupling by solid phase peptide synthesis ... 20
   2.7 Method for peptide acetylation ... 20
   2.8 Method for Alloc deprotection ... 21
   2.9 Method for TAMRA labeling ... 21
   2.10 Cleavage of peptides ... 21
   2.11 Purification of crude peptide ... 21
   2.12 Method for LC-MS characterization ... 21
   2.13 Analytical method for HPSEC-based affinity pull-down. ... 22
   2.14 Direct binding measurement by BLI ... 22
   2.15 Competitive binding assay by BLI ... 22
   2.16 Fluorescence polarization assay ... 23
   2.16 Cell culture ... 23
   2.17 Pull-down assay ... 23
   2.18 Imaging experiment with fixed cells ... 24
   2.19 Imaging experiments with live cells ... 24
   2.20 Transfection in HK-2 cells ... 25
3. LC-MS characterization of peptides ... 26
   3.1 Monomeric peptide ... 26
   3.2 Dimeric peptide ... 28
   3.3 Trimeric peptide ... 31
   3.4 Tetrameric peptide ... 33
4. References ... 35

### 1. Supplementary figures

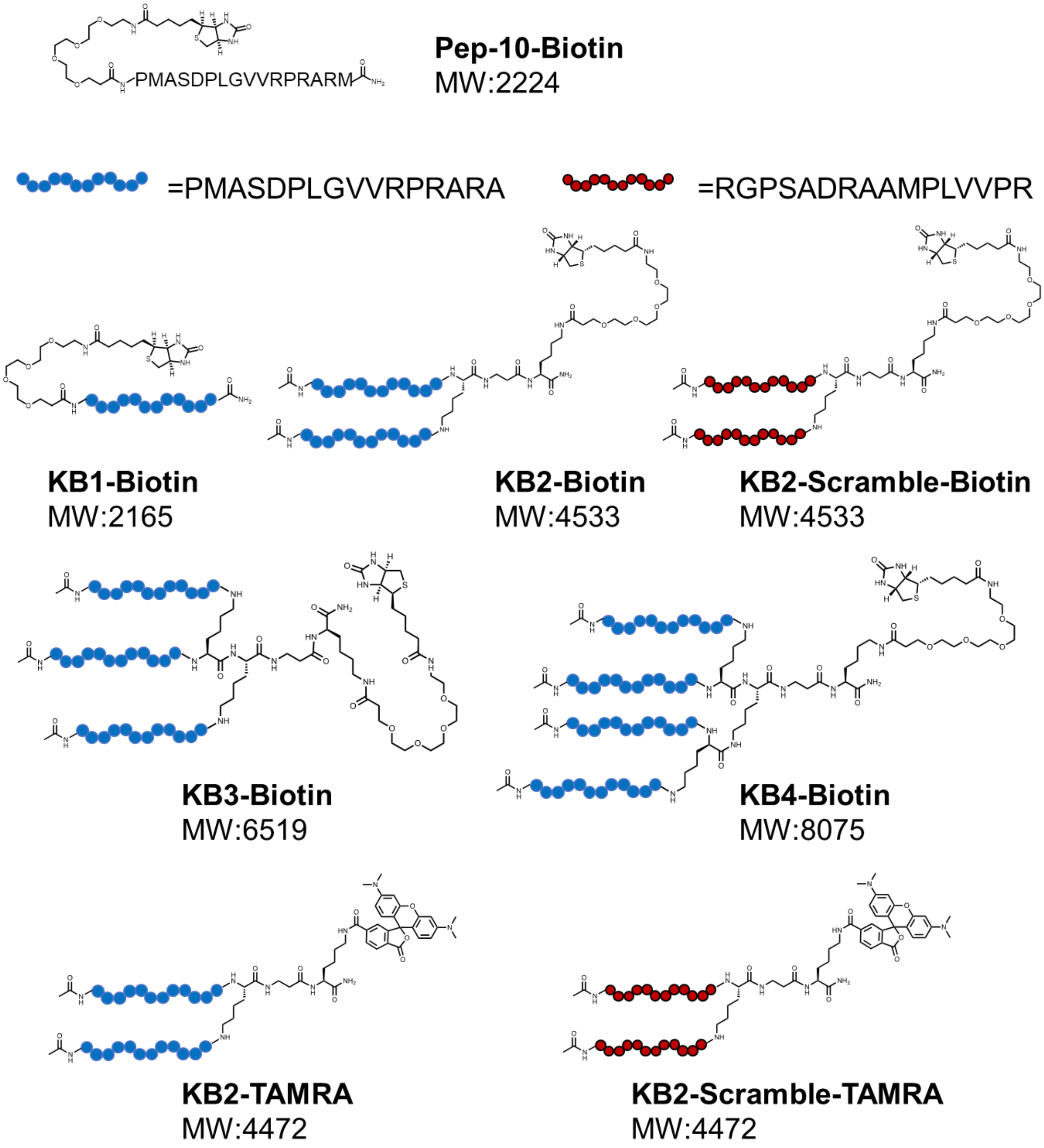


**Figure S1. Chemical structures of Pep-10-Biotin, KB1-Biotin, KB2-Biotin, KB2-Scramble-Biotin, KB3-Biotin, KB4-Biotin, KB2-TAMRA and KB2-Scramble-TAMRA.**

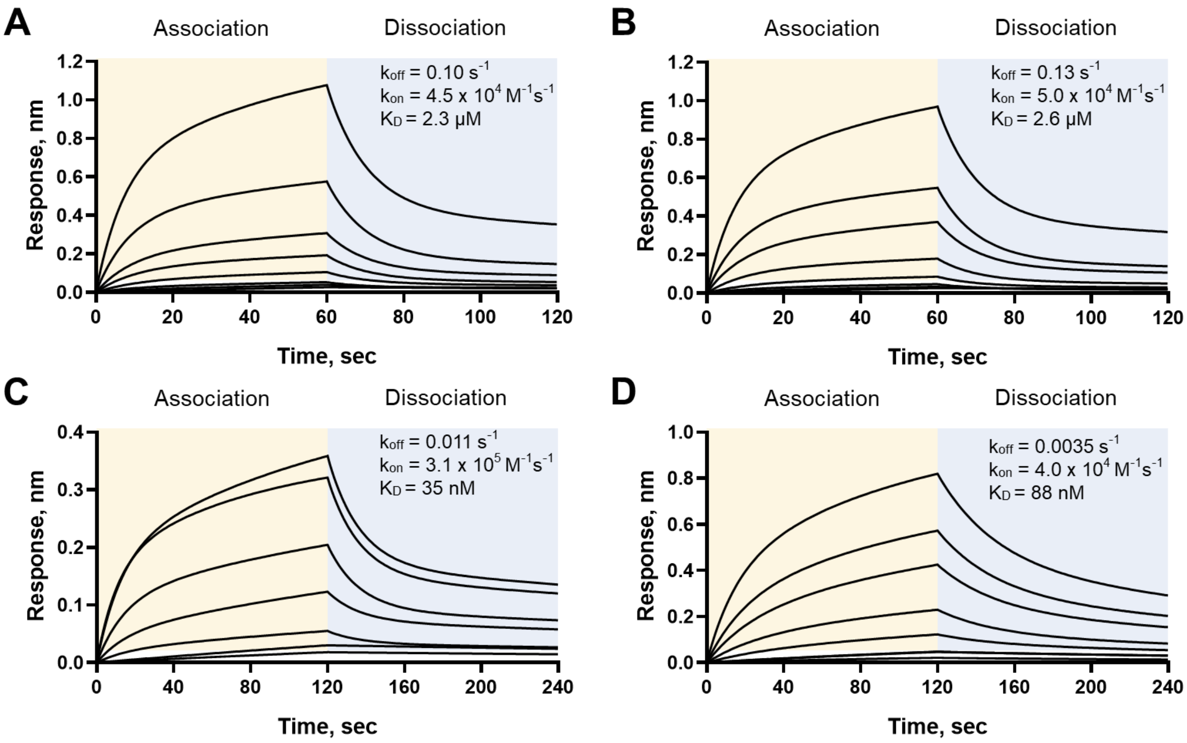


**Figure S2. A BLI-based direct binding analysis of Klotho-peptide interactions. Pep-10** (A), **KB1** (B), **KB3** (C), **KB4** (D) was functionalized with Biotin-PEG4 moiety using the preinstalled Alloc-handle. **Pep-10-Biotin** (A), **KB1-Biotin** (B), **KB3-Biotin** (C), **KB4-Biotin** (D) are immobilized onto streptavidin tips. The direct binding is recorded by a BLI system to study the dissociation and association kinetics (n = 2). The solutions contained various concentrations of Klotho as 1000, 500, 250, 125, 62.5, 31.3, 15.6, 7.8 nM.

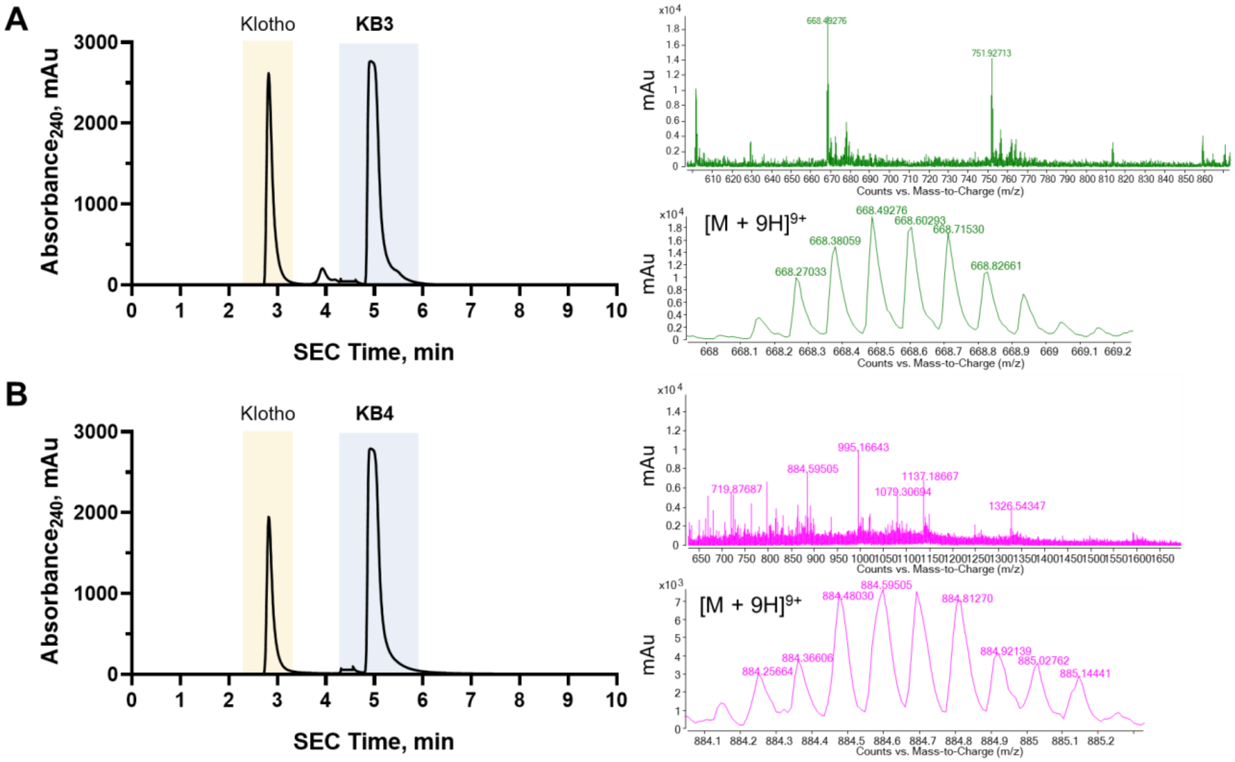


**Figure S3. The size-exclusion chromatography (SEC) analysis of Klotho–peptides complex.** SEC achieved a base-line separations of Klotho and peptides. SEC standard run was shown in **section 2.13**. MS of **KB3** (A) and **KB4** (B) were detected in the Klotho fractions (peak with retention time of 2.7 min) after precipitation of Klotho, indicating a strong Klotho-peptides interactions.

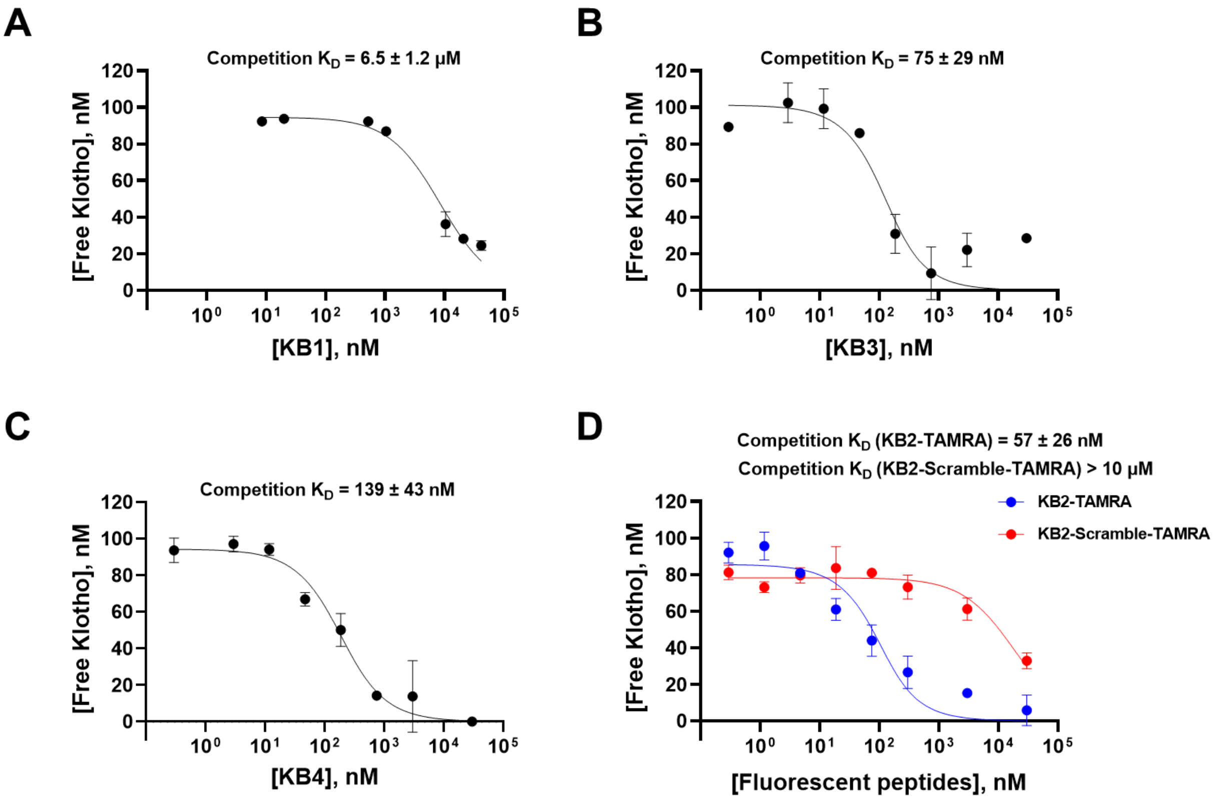


**Figure S4. The competition assay by BLI revealed compounds' competitive binding constants to Klotho.** (A) Unlabeled **KB1** binds to mouse Klotho (mKlotho) weakly with estimated competitive $K_D$ over 5000 nM. (B) Unlabeled **KB3** binds to mKlotho with competitive $K_D$ = 75 ± 29 nM. (C) Unlabeled **KB4** binds to mKlotho with competitive $K_D$ = 139 ± 43 nM. (D) **KB2-TAMRA** binds to mKlotho with competitive $K_D$ = 57 ± 26 nM. **KB2-TAMRA**'s competitive binding constant is similar to **KB2** in this system. All assays are done in duplicate. Error bars indicate SD.

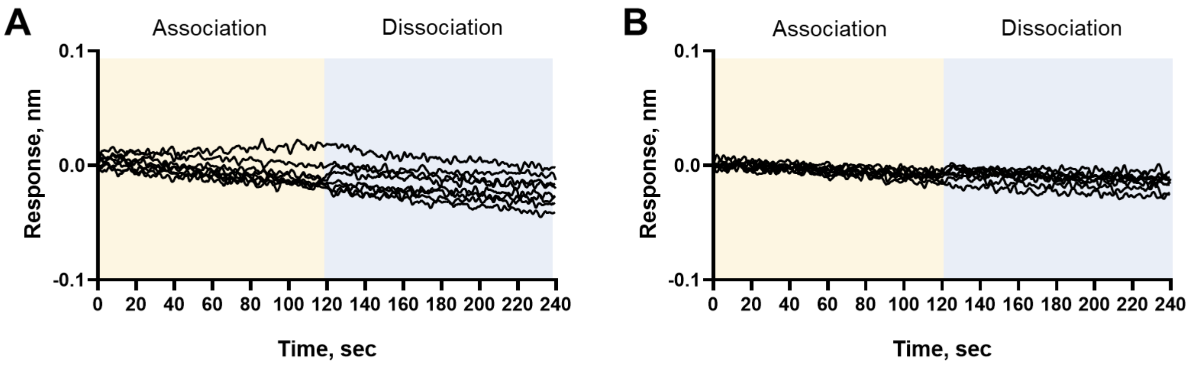


**Figure S5. A BLI-based binding analysis of (A) MDM2 and (B)12CA5-peptide interactions. KB2-Biotin** is immobilized onto streptavidin tips. No binding was observed between **KB2-Biotin** and these two proteins (n = 2). The solutions contained various concentrations of (A) MDM2 and (B) 12CA5 as 1000, 500, 250, 125, 62.5, 31.3, 15.6, 7.8 nM.

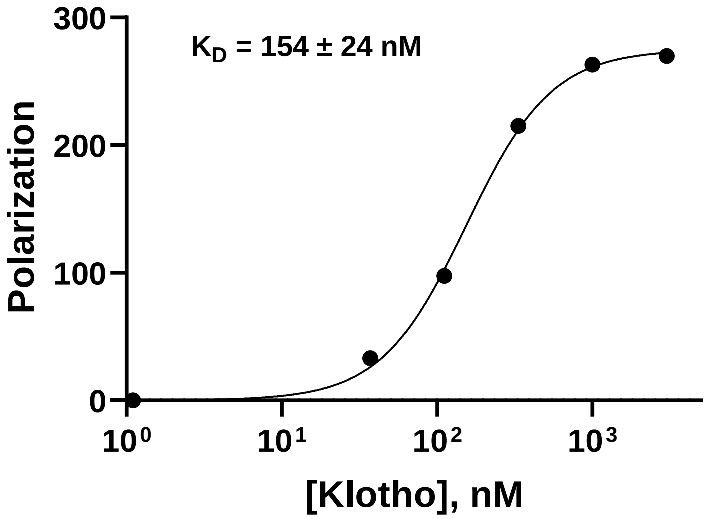


**Figure S6. Fluorescence polarization assay was established to estimate the binding dissociation constant of KB2-TAMRA and Klotho.** Klotho (3 μM) was prepared in 1× PBS followed by a 1:2 serial dilution supplemented with **KB2-TAMRA** to a final concentration of 100 nM. Polarization was measured with excitation wavelength of 545 nm and emission wavelength of 595 nm (n = 2). Error bars indicate SD.

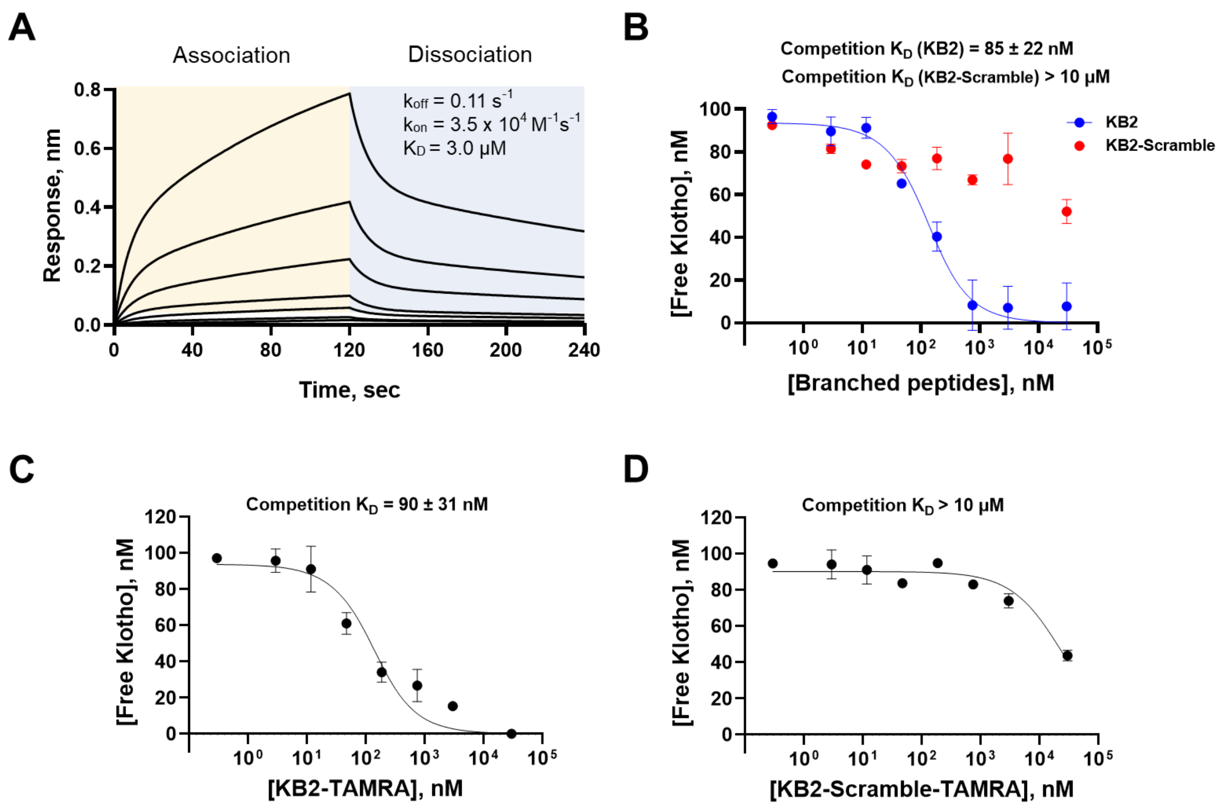


**Figure S7. A BLI-based binding analysis of hKlotho-peptide interactions.** (A) **KB1** with Biotin-PEG4 moiety using the preinstalled Alloc-handle. **KB1-Biotin** is immobilized onto streptavidin tips. The direct binding is recorded by a BLI system to study the dissociation and association kinetics (n = 2). (B) The competition assay by BLI revealed compounds' competitive binding constants against hKlotho. Unlabeled **KB2** binds to hKlotho with competitive $K_D$ = 85 ± 22 nM while **KB2-Scramble** has minimal binding. (C) **KB2-TAMRA** binds to hKlotho with competitive $K_D$ = 90 ± 31 nM. **KB2-TAMRA**'s competitive binding constant is similar to **KB2** against hKlotho. (D) **KB2-Scramble** has minimal binding to hKlotho with competitive $K_D$ over 10 μM. All assays are done in duplicate. Error bars indicate SD.

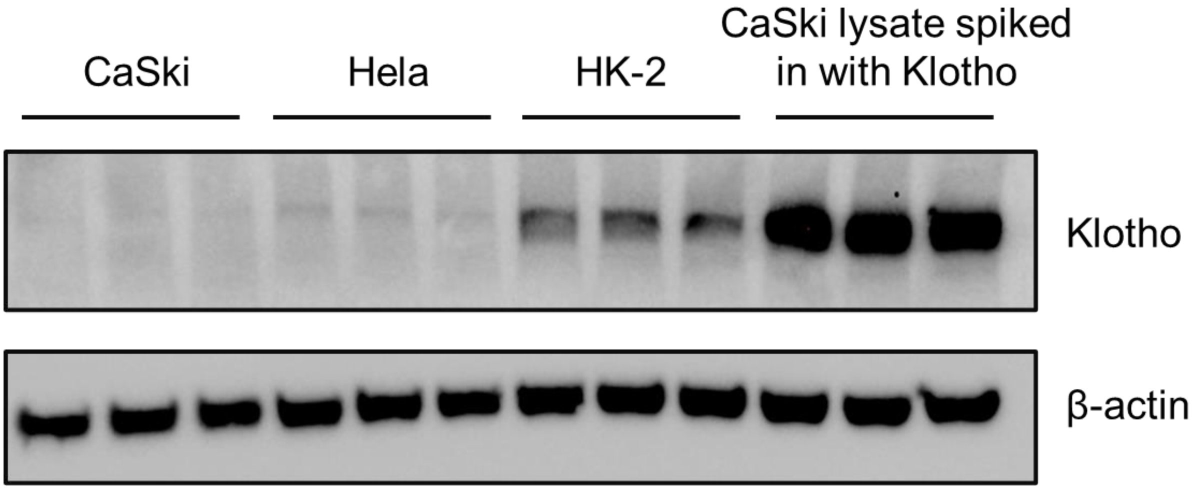


**Figure S8. Western blot gels demonstrated Klotho level in different cell lines.** The first three conditions are 100 μg of CaSki, Hela and HK-2 cells. The fourth condition is 100 μg of CaSki lysate spiked in with ~0.5 μg of recombinant mKlotho (final concentration is 0.1 μM in 50 μL).

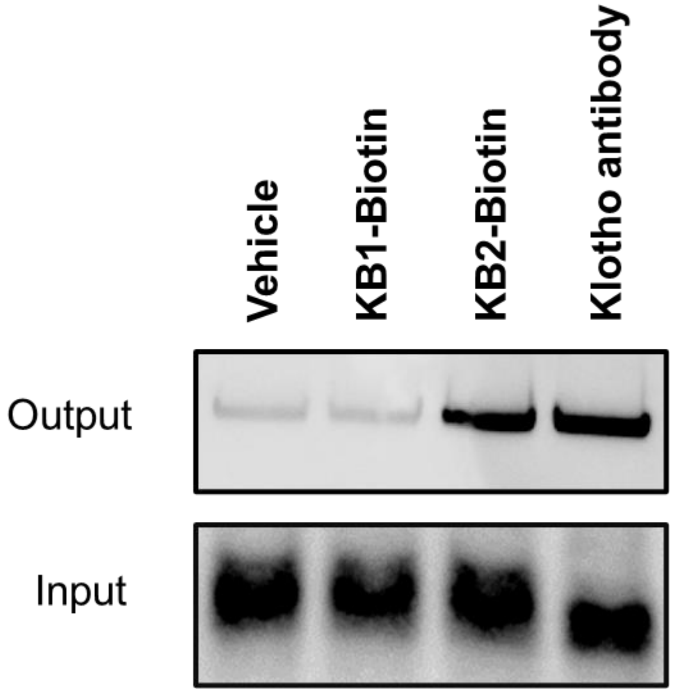


**Figure S9. Pull-down assays from human cell lysates.** Pull-down probe **KB2-Biotin** at 200 nM enriched recombinant mKlotho from 100 µg of CaSki cell lysate while **KB1-Biotin** displayed no enrichment of Klotho at 1 µM. Pull-down by Klotho antibody (KM2076) was performed using Dynabeads™ Sheep Anti-Rat IgG (Invitrogen, 11035). Pull-down fraction of Klotho antibody gave bands with the same molecular weight, confirming it was Klotho being enriched by biotinylated peptides.

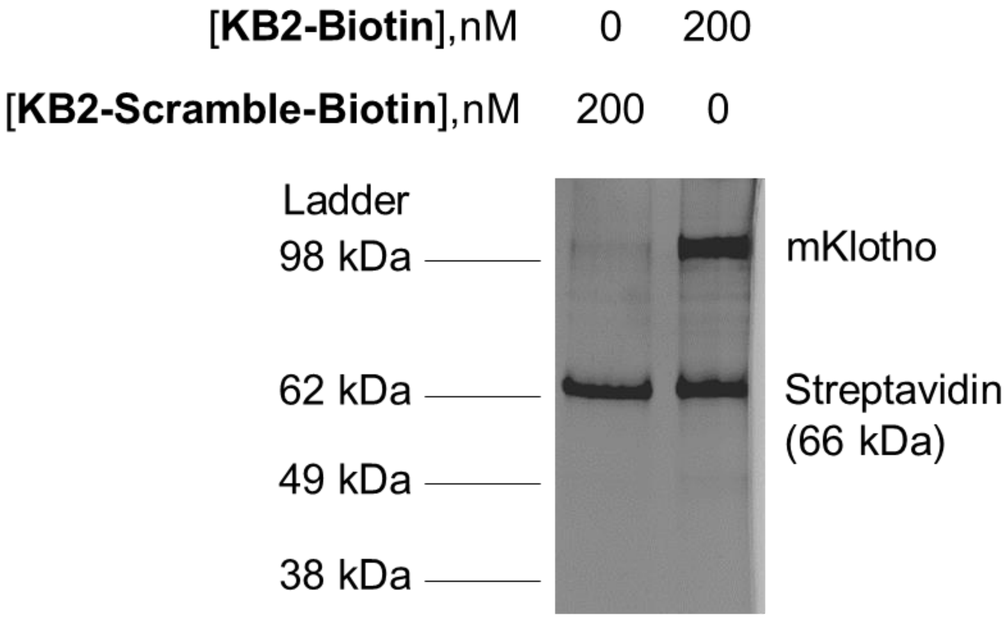


**Figure S10. Silver stain analysis of pull-down from CaSki lysate spiked in with mKlotho (0.1 µM).** Pull-down probe **KB2-Biotin** at 200 nM enriched recombinant mKlotho from 100 µg of CaSki cell lysate at 200 nM. Control probe **KB2-Scramble-Biotin** displayed negligible enrichment of Klotho at the same concentration. Other protein pull-down bands are similar across two lanes, which indicated selective pull-down of Klotho over other proteins in lysate.

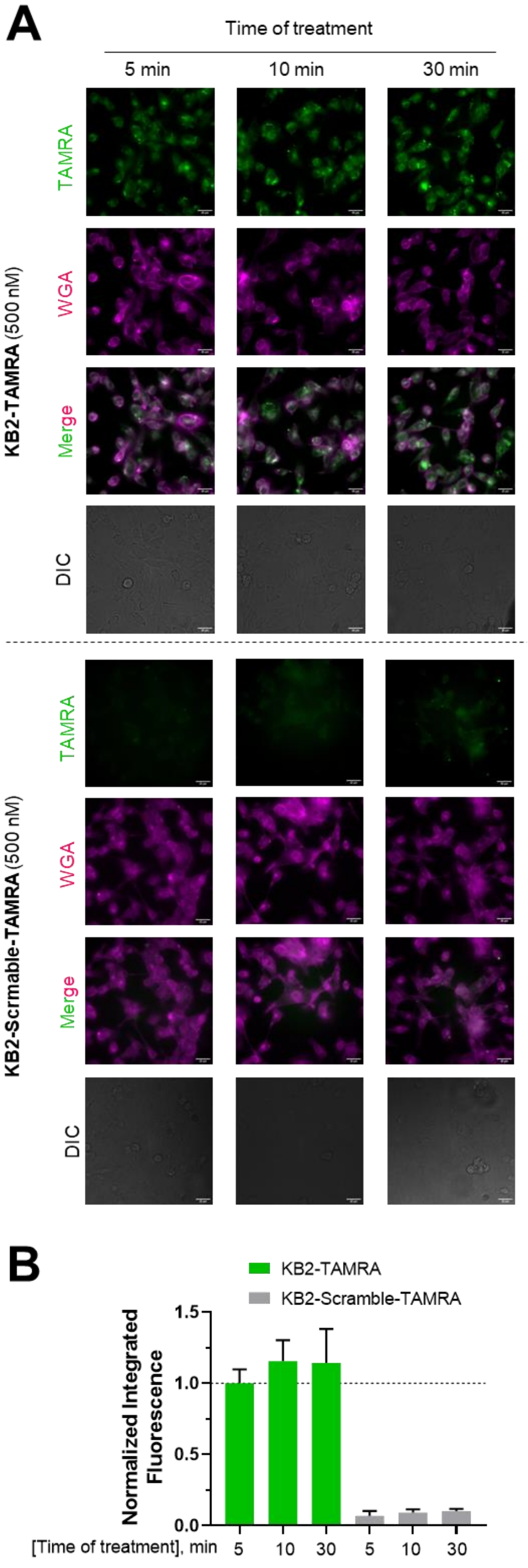
A
Time of treatment
5 min
10 min
30 min
KB2-TAMRA (500 nM)
TAMRA
WGA
Merge
DIC
KB2-Scrmable-TAMRA (500 nM)
TAMRA
WGA
Merge
DIC
B
KB2-TAMRA
KB2-Scramble-TAMRA
Normalized Integrated Fluorescence
1.5
1.0
0.5
0.0
[Time of treatment], min
5
10
30
5
10
30

**Figure S11. Time course study of labeling of Klotho in fixed HK-2 cells by fluorescence microscopy imaging.** (A) Three different time of treatment (5, 10, 30 min) were used. **KB2-TAMRA** (top) and **KB2-Scramble-TAMRA** (bottom) at 500 nM was co-treated with Wheat Germ Agglutinin (WGA) Alexa Fluor™ 350 Conjugate at 25 ng/µL in HK-2 cells. Fixation of cells was performed after the treatment. Fluorescence of fluorescent peptides was acquired in the orange channel ($\lambda_{ex/em}$ = 542/597 nm). Fluorescence of WGA Alexa Fluor™ 350 was acquired in the blue channel ($\lambda_{ex/em}$ = 390/435 nm). Scale bar = 25 µm. (B) The bar graph depicts quantification of normalized integrated fluorescence (n = 10). Error bars indicate SD.

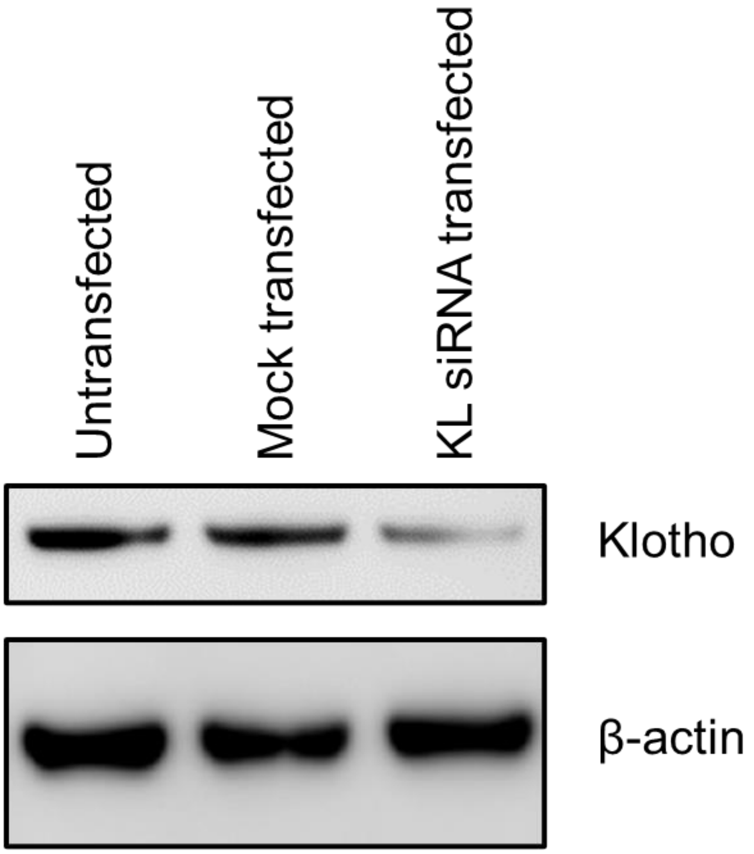


**Figure S12. Representative western blot gels demonstrated KL siRNA downregulates Klotho level in HK-2 cells. n = 2.**

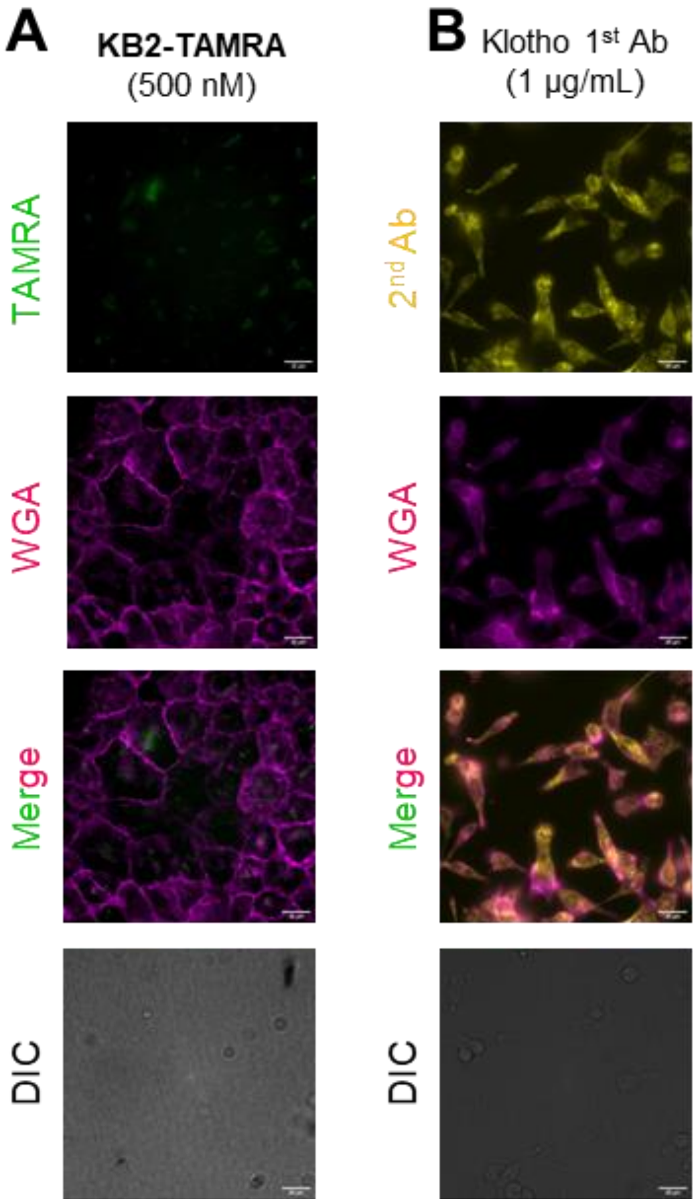


**Figure S13. Characterization of KB2-TAMRA revealed selective labeling of Klotho.** (A) Detection of intracellular fluorescence and potential labeling of endogenous total hKlotho following 5 min of co-treatment with **KB2-TAMRA** at 500 nM and Wheat Germ Agglutinin (WGA) Alexa Fluor™ 350 Conjugate at 25 ng/µL in CaSki cells. Fixation of cells was performed after the treatment. **KB2-TAMRA** does not display appreciable fluorescence in CaSki cells with low levels of hKlotho expression. Fluorescence of **KB2-TAMRA** was acquired in the orange channel ($\lambda_{ex/em}$ = 542/597 nm). Fluorescence of WGA Alexa Fluor™ 350 was acquired in the blue channel ($\lambda_{ex/em}$ = 390/435 nm). (B) Labeling of endogenous total hKlotho by overnight incubation of primary Klotho antibody (1st Ab) and 1 h incubation of secondary antibody (2nd Ab) Goat anti-Rat IgG Alexa Fluor™ 647 in fixed HK-2 cells. Fluorescence of 2nd Ab was acquired in the red channel ($\lambda_{ex/em}$ = 632/679 nm). Scale bar = 25 µm.

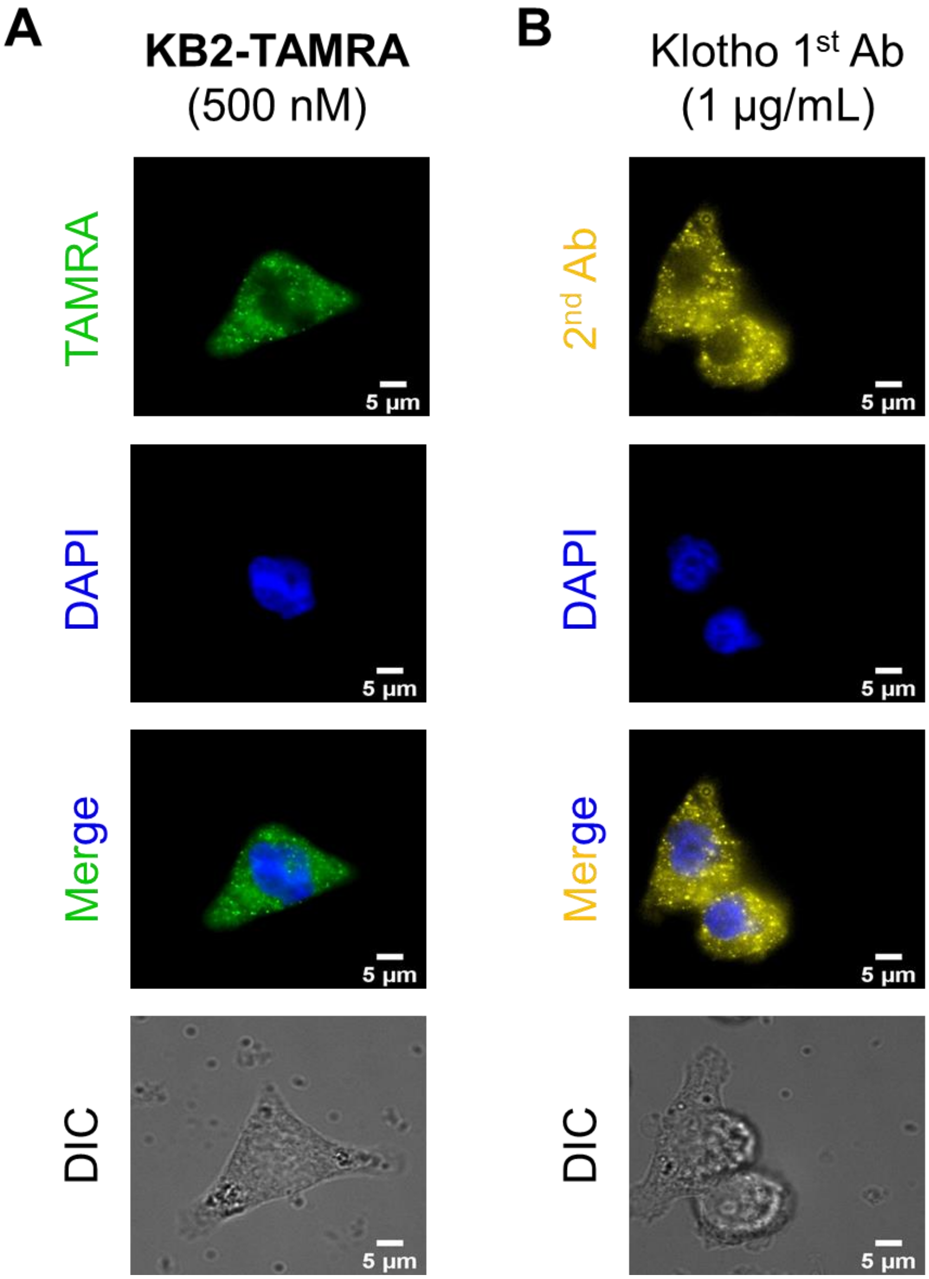


**Figure S14. Imaging experiments showed distribution of hKlotho in HK-2 cells.** (A) Labeling of endogenous hKlotho by 5 min of treatment of **KB2-TAMRA** at 500 nM in HK-2 cells. Fixation of cells was performed after the treatment. After fixation, cells were stained with DAPI (300 nM) for 5 min. Fluorescence of **KB2-TAMRA** was acquired at orange channel (Ex/Em = 542/597 nm). Fluorescence of DAPI was acquired at blue channel (Ex/Em = 390/435 nm). (B) Labeling of endogenous hKlotho by overnight incubation of primary antibody ($1^{st}$ Ab) of Klotho and 1 h incubation of secondary antibody ($2^{nd}$ Ab) Goat anti-Rat IgG Alexa Fluor 647 in fixed HK-2 cells. Fluorescence of $2^{nd}$ Ab was acquired at red channel (Ex/Em = 632/679 nm). After incubation with $2^{nd}$ Ab, cells were stained with DAPI (300 nM) for 5 min. Pearson correlation coefficients were used to quantify co-localization of nucleus dye DAPI versus **KB2-TAMRA** and Klotho antibody. The coefficient between DAPI and **KB2-TAMRA** is 0.29. The coefficient between DAPI and Klotho antibody is 0.35. Scale bar = 5 μm.

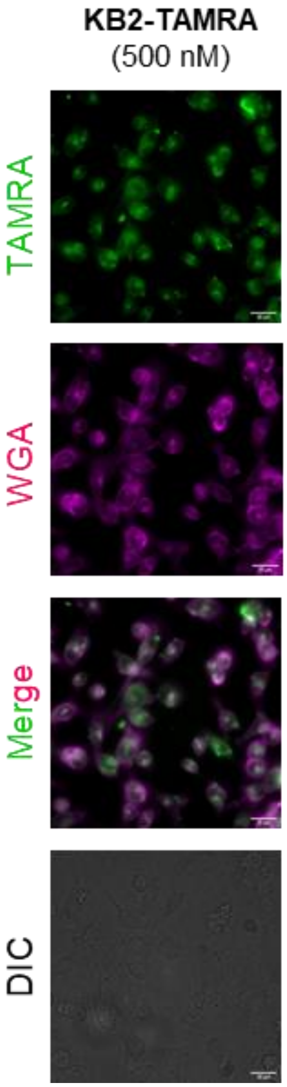


**Figure S15. Fluorescence microscopy imaging revealed KB2-TAMRA labeling total Klotho in live HK-2 cells.** Detection of intracellular fluorescence and potential labeling of endogenous total hKlotho following 5 min of co-treatment with **KB2-TAMRA** at 500 nM and WGA Alexa Fluor™ 350 Conjugate at 25 ng/µL in *live* HK-2 cells. Images were taken after the treatment without fixation. Scale bar = 25 µm.

### 2. Methods and Materials

#### 2.1 Materials.

H-Rink Amide-ChemMatrix resin was purchased from PCAS BioMatrix Inc. Amino acids: Fmoc-Ala-OH, Fmoc-β-Ala-OH, Fmoc-Asn(Trt)-OH, Fmoc-Asp(*t*Bu)-OH, Fmoc-Cys(Trt)-OH, Fmoc-Glu(*t*Bu)-OH, Fmoc-Leu-OH, Fmoc-Lys(Boc)-OH, Fmoc-Phe-OH, Fmoc-Pro-OH, Fmoc-Ser(*t*Bu)-OH, Fmoc-Thr(*t*Bu)-OH, Fmoc-Trp(Boc)-OH, Fmoc-Tyr(*t*Bu)-OH, and Fmoc-Val-OH were purchased from Novabiochem (Billerica, MA). Other amino acids: Fmoc-Lys(ivdde)-OH, Fmoc-Lys(Alloc)-OH, Fmoc-Lys(Fmoc)-OH were purchased from Novabiochem(Billerica, MA). Palladium tetrakis (Pd(PPh3)4) was purchased from Sigma Aldrich. Reagents used in solid phase peptide synthesis**:** ChemMatrix H-Rink Amide (0.18 mmol/g loading) resin was purchased from PCAS Biomatrix. Piperidine (ReagentPlus; 99%), formic acid (≥98%) were purchased from Sigma-Aldrich (St. Louis, MO). Diisopropylethylamine (DIEA; biotech. grade; 99.5%) was purchased from MilliporeSigma and purified by a Seca Solvent Purification system from Pure Process Technology (Nashus, NH). Reagents (cleavage): Trifluoroacetic acid (TFA; for HPLC, ≥99%), triisopropylsilane (TIPS; 98%), were purchased from Sigma-Aldrich (St. Louis, MO). Reagents used in peptide post-synthesis modifications: Acetic anhydride (≥98%) was purchased from Sigma-Aldrich (St. Louis, MO). 5-TAMRA (5-carboxytetramethylrhodamine) and Biotin-PEG4-carboxylic acid from ChemPep. Human Klotho protein was ordered from R&D Systems (5334-KL-025). MDM2 protein was ordered from Abcam (ab167941). 12CA5 was ordered from Columbia Biosciences (00-1722). KL siRNA (SMARTpool: ON-TARGETplus Human KL siRNA) was purchased from horizon with catalog number (L-011936-01-0005).

Cell lines, CaSki (ATCC CRM-CRL-1550), HK-2 (ATCC CRL-2190), were obtained from ATCC. Media (RPMI-1640 and Keratinocyte SFM), fetal bovine serum (FBS), penicillin- streptomycin, X10 PBS and 0.25% trypsin-EDTA were obtained from Gibco. Culture flasks, plates, and serological pipettes were obtained from Fisher scientific. Fixing reagent: 37% formaldehyde, dilute 10 time in 1x PBS (sigma). Dye: Wheat Germ Agglutinin (WGA) Alexa Fluor™ 350 Conjugate (W11263, Thermo) and DAPI (Invitrogen).

#### 2.2 Mouse Klotho (mKlotho) protein production

Mouse Klotho(aa1-960)-aaa-6His was produced from a HEK-293-H stable cell line (Thermo Fisher Scientific Inc. Cat: 11631017) in Hybridoma-SFM (Thermo-Fisher) media supplemented with 2% FBS-HI, 1% HEPES, and 1% PenStrep. After seven days, 10 liters of conditioned media was centrifuged and filtered and applied to a 5 mL Histrap Excel column (5ml bed volume, Cytiva 17-3712-05). The Histrap Excel column was washed with 50 mM HEPES pH 7.5, 150 mM NaCl, 50 mM imidazole and eluted with 50 mM HEPES pH 7.5, 150 mM NaCl, 500 mM imidazole. mKlotho was then polished over a HiLoad Superdex S200 PG Column (Cytiva 28989336). Purified protein fractions from size exclusion chromatography were concentrated to approximately 1 mg/ml in 1X PBS and stored at -80°C for further analysis.

#### 2.3 Fast flow synthesis of branched peptides

H-Rink Amide-ChemMatrix resin (200 mg, 0.49 mmol/g, 0.10 mmol) was used to prepare peptides. Peptides were prepared by fully automated SPPS[1]. Upon completion, resins were washed with DCM (3x) and dried under reduced pressure.

#### 2.4 Manual preparation of peptidyl resins with linker 1,2, and 3

ChemMatrix® Rink amide resin (loading 0.18 mmol/g, typical scale: 100 mg, 0.02 mmol) was loaded into a fritted syringe (6 mL), swollen in DMF (4 mL) for 5 minutes and then drained. Each Nα-Fmoc protected amino acid (0.2 mmol, 10 equiv) was dissolved in DMF containing 0.39 M HATU (0.5 mL). Immediately before the coupling DIEA (100 µL, 30 equiv) was added to the mixture to activate the amino acid. After 15 seconds pre-activation the mixture was added to the resin and reacted for 10 min, with occasional stirring. After completion of the coupling step, the syringe was drained, and the resin was washed with DMF (3 x 5 mL). Fmoc deprotection was performed by addition of piperidine (20% in DMF, 3 mL), to the resin (2 x 5 min), followed by draining and washing the resin with DMF (5 x 5 mL). ivdde deprotection was performed by addition of hydrazine (3% in DMF, 3mL), for 1 min. For dimer peptidyl resin linker 1 the coupling cycles were performed sequentially with Fmoc-Lys(Alloc)-OH, Fmoc-β-Ala-OH and Fmoc- Lys(Fmoc)-OH; for trimer peptidyl resin with linker 2 the coupling cycles were performed sequentially with Fmoc- Lys(Alloc)-OH, Fmoc-β-Ala-OH, Fmoc-Lys(ivdde)-OH, deprotect ivdde and couple Fmoc-Lys(Fmoc)-OH; for tetramer peptidyl resin with linker 3 the coupling cycles were performed sequentially with Fmoc- Lys(Alloc)-OH, Fmoc-β-Ala-OH, Fmoc-Lys(Fmoc)-OH and Fmoc-Lys(Fmoc)-OH.

#### 2.5 Automated flow peptide synthesis (AFPS) set-up

All peptides were synthesized on automated-flow systems built in the Pentelute lab ("Amidator" and "Peptidator"), which are similar to the published AFPS system. The synthesis conditions used are according to Hartrampf [2]et al.:

The following settings were used for protein synthesis: flowrate = 40 mL/min, temperature = 90 °C (loop) and 85–90 °C (reactor). The 50 ml/min pump head pumps 400 μL of liquid per pump stroke; the 5 mL/min pump head pumps 40 μL of liquid per pump stroke. The standard synthetic cycle involves a first step of prewashing the resin at elevated temperatures for 60 s at 40 mL/min. During the coupling step, three HPLC pumps are used: a 50 mL/min pump head pumps the activating agent, a second 50 ml/min pump head pumps the amino acid and a 5 mL/min pump head pumps DIEA. The first two pumps are activated for 8 pumping strokes in order to prime the coupling agent and amino acid before the DIEA pump is activated. The three pumps are then actuated together for a period of 7 pumping strokes, after which the activating agent pump and amino acid pump are switched using a rotary valve to select DMF. The three pumps are actuated together for a final 8 pumping strokes, after which the DIEA pump is shut off and the other two pumps continue to wash the resin for another 40 pump strokes. During the deprotection step, two HPLC pumps are used. Using a rotary valve, one HPLC pump selects deprotection stock solution and DMF. The pumps are activated for 13 pump strokes. Both solutions are mixed in a 1:1 ratio. Next, the rotary valves select DMF for both HPLC pumps, and the resin is washed for an additional 40 pump strokes. The coupling–deprotection cycle is repeated for all additional monomers.

#### 2.6 Linker coupling by solid phase peptide synthesis

200 mg of H-Rink Amide-ChemMatrix resin (0.10 mmol) was placed in a 5 mL Torviq fritted syringe. After swelling in DMF for 10 min, the resin was then washed with DMF (3x). Each Fmoc-protected amino acid (0.50 mmol, 5.0 eq) was dissolved in 0.38 M HATU solution (DMF as solvent, 1.3 mL, 0.50 mmol). Immediately before the coupling, DIEA (130 μL, 0.72 mmol, 6.9 eq) was added to the mixture. The resulting mixture was sonicated briefly, and then transferred to the fritted syringe containing the resin. Coupling was performed for 20 min. The resin was washed with DMF (3x). Fmoc deprotection was done by treating the resin with 20% piperidine in DMF (3 mL) for 5 min and this step was repeated twice. The resulting resin was then washed with DMF (3x). The synthetic cycle was repeated to completion of the peptide sequence.

#### 2.7 Method for peptide acetylation

A 100 mg portion of peptidyl resin from section 6.1.1 was placed into a 5 mL Torviq fritted syringe and subsequently swelled in DMF. After removing DMF, a solution of Ac2O, DIEA, and DMF (2 mL, 85:315:1600, v/v) was added to the peptidyl resin. The resulting mixture was occasionally agitated for 45 min. After draining the solution, the remaining resin was washed with DMF (3x), DCM (3x) and dried under reduced pressure.

#### 2.8 Method for Alloc deprotection

Peptidyl resin (~10 μmol theoretical loading) was washed with dichloromethane (3 x 5 mL) and then treated with Pd(PPh3)4 (11.0 mg, 10 μmol, 1 equiv) in dichloromethane/piperidine (8:2, 1 mL) for 30 minutes at room temperature under exclusion of light. The resin was then drained and washed with dichloromethane (3 x 5 mL).

#### 2.9 Method for TAMRA labeling

Peptidyl resin (~ 10 μmol theoretical loading) was loaded into a fritted syringe (6 mL), swollen in DMF (4 mL) for 5 minutes and then drained. 5-Carboxytetramethylrhodamine (5-TAMRA, 22 mg, 50 μmol, 5 equivalents) and HATU (17 mg, 45 μmol, 4.5 equivalents) were dissolved in DMF (500 μL), activated with DIEA (19 mg, 26 μL, 150 μmol), added to the peptidyl resin and incubated for 30 minutes under exclusion of light. After this time the resin was drained, washed with DMF (3 x 5 mL) and stored until cleavage.

#### 2.10 Cleavage of peptides

Synthesized peptide was cleaved from resin and globally deprotected by treating the peptidyl resin with a cleavage cocktail containing 94% TFA, 2.5% EDT, 2.5% water, and 1.0% TIPS (*v/v*), for 1 h at ambient temperature. TFA was removed under a gentle stream of nitrogen gas, and crude peptide was precipitated by the addition of cold $Et_2O$ (-80 °C). After centrifugation at 3220 rcf for 3 min, supernatant was removed, and precipitated peptide was triturated three times with cold $Et_2O$. The resulting material was dissolved in 50% MeCN in water with 0.1% TFA, and lyophilized.

#### 2.11 Purification of crude peptide

Crude peptides were purified by a Biotage Selekt flash purification system. Water with 0.1% TFA (solvent A) and MeCN with 0.1% TFA (solvent B) were utilized as mobile phases for purifications. The crude peptide was dissolved in minimal amount of 10% MeCN in water with 0.1% TFA, and then loaded onto a 10 g Biotage SNAP Bio C4 20 μm column. The purification was performed using a gradient as following: 10% B for 2 column volume (CV), linear ramp from 30% B to 50% B for 20 CV, 25 mL/min flow rate.

#### 2.12 Method for LC-MS characterization

LC-MS characterizations were carried out using an Agilent 6550 quadrupole time-of-flight LC-MS. Total ion current (TIC) chromatograms were plotted. Mass spectra were integrated over the principal TIC peaks. High-performance liquid chromatography was done by following methods: (solvent A: water with 0.1% formic acid; solvent B: MeCN with 0.1% formic acid). **Method A:** Column: Phenomenex Jupiter C4 column (1.0 x 150 mm, 5 μm particle size, 300 Å pore size) Gradient: 1% B (0-2 min), linearly ramp from 1% B to 91% B (2-8 min). Flow rate is 100 μL/min. MS acquisition is from 2 to 8 min. **Method B:** Column: Phenomenex Jupiter C4 column (1.0 x 150 mm, 5 μm particle size, 300 Å pore size) Gradient: 1% B (0-2 min), linearly ramp from 1% B to 61% B (2-12 min), 61% B to 95% B (11-16 min). Flow rate is 100 μL/min. MS acquisition is from 4 to 12 min. **Method C:** Column: Agilent Zorbax 300SB C3 column (2.1 x 150 mm, 5 μm particle size, 300 Å pore size) Gradient: 1% B (0-2 min), linearly ramp from 1% B to 91% B (2-12 min), 91% B to 91% B (12-13 min). Flow rate is 500 μL/min. MS acquisition is from 4 to 12 min.

#### 2.13 Analytical method for HPSEC-based affinity pull-down.

High-performance size exclusion chromatography (HPSEC) was carried out using an Agilent 1260 Infinity II LC System with the Agilent BIOSEC-3 HPLC column (7.8 x 150 mm, 3 μm particle size, 100 Å pore size). HPSEC samples, such as proteins, peptide, or protein-peptide mixtures (5 min incubation at 4 °C), were prepared in 100 μL 1x PBS buffer, and then eluted in buffered mobile phase at 1 mL/min flow rate for 15 min. While performing affinity selection experiments, the protein-binder complex fraction was detected by UV (214 and 280 nm) and collected for LC-MS analysis using Method A in the above LC-MS method session. Peptide recovery was calculated based on the extracted ion count integration peak area ratio of peptide solution before and after the HPSEC separation. After each HPSEC affinity selection experiment, SEC column was cleaned with an IPA/water/MeCN/MeOH (1:1:1:1, *v/v*) mixture containing 0.1% formic acid (FA).

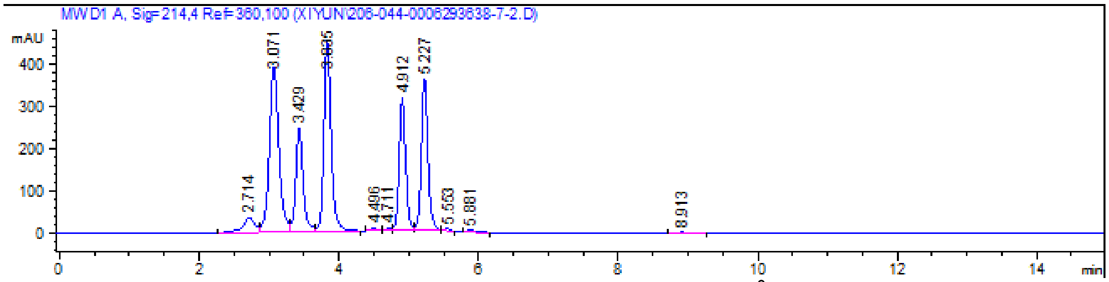


SEC standard run with Agilent™Agilent BioSEC standard 130Å (Agilent, MA). Buffer: 150 mM $Na_3PO_4$, pH = 7. Flow rate: 1 mL/min. Five peaks correspond to Ovalbumin, 45 kDa; Myoglobin 17 kDa; Aprotinin, 6.7 kDa; Neurotensin, 1.7 kDa; Angiotensin II, 1.0 kDa.

#### 2.14 Direct binding measurement by BLI

Streptavidin sensors were soaked in blocking buffer (PBS supplemented with 0.05% Tween-20 and 1 mg/mL bovine serum albumin) for 5 min. After immobilizing the biotinylated peptides (200 nM) onto streptavidin sensors, 2-time serial dilutions of Klotho in blocking buffer were analyzed for binding. The solutions contained various concentrations of Klotho as 1000, 500, 250, 125, 62.5, 31.3, 15.6, 7.8 nM. The response was recorded after 2 min when BLI signal reaches equilibrium. Association lasted for 120 sec (60 sec for **KB1-Biotin**) and dissociation last for another 120 sec (60 sec for **KB1-Biotin**). The curve was reported by GatorPlus software and replotted by Prism 8 software. Error bars of $K_D$ reported in the main body indicate SD.

#### 2.15 Competitive binding assay by BLI

A competition binding assay was performed as described below using GatorPlus bio-layer interferometry(GatorBio) to estimate the binding affinity of peptides.

*Calibration curve:* Streptavidin sensors were soaked in blocking buffer (PBS supplemented with 0.05% Tween-20 and 1 mg/mL bovine serum albumin) for 5 min. After immobilizing the PEG4-Biotinylated **KB1** peptide (200 nM of Biotin-PEG4-PMASDPLGVVRPRAA) onto streptavidin sensors, serial dilutions of Klotho in blocking buffer were analyzed for binding. The solutions contained various concentrations of Klotho as 1000, 500, 250, 125, 62.5, 31.3, 15.6, 7.8 nM. The response was recorded after 2 min when BLI signal reaches equilibrium. A curve of sensor response (nm) vs. Klotho concentration (nM) was generated to calibrate the free Klotho concentration in solution observed in the competition assay. The curve was generated using Prism 8.

*Competition assay.* Various concentrations of peptides were incubated in wells with Klotho protein in the blocking buffer for 30 min. The PEG4-Biotinylated **KB1** peptide was immobilized onto streptavidin sensors and dipped into preincubated sample wells. The association events were measured at 30 °C, 1,000 rpm. The response was recorded after 2 min when BLI signal reaches equilibrium. Based on the binding response (nm) values, the concentration of 'free' Klotho was interpolated for each sample using the calibration curve. The apparent dissociation constant, $K_D$, can be obtained from the non-linear regression analysis using the equation $[\mathrm{Y}] = 0.5 \times [b - \mathrm{K}_d - [\mathrm{X}] + \sqrt{([\mathrm{X}] + \mathrm{K}_d - b)^2 + 4b \times \mathrm{K}_d}\ ]$ ,where [Y] is the free [Klotho] in nM, [X] is the total [peptide] in nM, $K_D$ is the binding dissociation constant to be fitted by the equation, and b is the maximal possible Klotho concentration to be fitted by the equation. By fitting the free [Klotho] and [peptide] to the equation, a binding constant with a fitting error were generated by Prism 8 software.

### 2.16 Fluorescence polarization assay

Klotho (3 µM) was prepared in 1× PBS followed by a 1:2 serial dilution. The protein solutions were then supplemented with **KB2-TAMRA** to a final concentration of 100 nM. The solutions were incubated at room temperature for 30 min in the dark. Polarization was measured using a Tecan plate reader with excitation wavelength of 545 nm and emission wavelength of 595 nm.

### 2.16 Cell culture

Cells were cultured at 37 °C with 5% $CO_2$. CaSki cells (ATCC, CRL-1550) were cultured in RPMI 1640 medium (ATCC, 30-2001) supplemented with 1× Pen Strep solution (Gibco, 15140-122) and 10% fetal bovine serum (FBS; Sigma, 12003C). HK-2 cells (ATCC, CRL-2190) was cultured in Keratinocyte SFM (Gibco, 17005042) supplemented with bovine pituitary extract and human recombinant EGF.

### 2.17 Pull-down assay

CaSki cells were washed with 1× PBS and harvested by trypsinization. Total protein was extracted with RIPA lysis buffer (Millipore, 20-188) per the manufacturer's protocol. Recombinant mouse Klotho (mKlotho) was added into 100 µg of CaSki cell lysate to final concentration of 0.1 µM. **KB2-Biotin** or **KB2-Scramble-Biotin** was then added into the protein solution and incubated at room temperature for 30 min. For the competitive pull-down assay, unlabeled **KB2** and **KB2-Scramble** was pre-incubated in the protein solution for 15 min before addition of biotinylated probes. After incubation, streptavidin magnetic beads (Dynabeads MyOne Streptavidin T1, Invitrogen, 65601) was added into the solution to pull down Klotho, and samples were shaken at room temperature for 20 min. The samples were washed with 1× PBS. The proteins were eluted from the beads by 30 µL of 8M urea solution from each sample then resolved on an SDS-polyacrylamide gel (Bolt 4-12% Bis-Tris Plus, Invitrogen, NW04120BOX), followed by protein transfer to a nitrocellulose (0.45 µm) membrane. After blocking the membrane in 1× TBST with 5% (w/v) nonfat milk for 1 h, the membrane was incubated with 1× TBST containing 5% milk and Klotho (Cosmo Bio, KM2076) primary antibody (1:1000 dilution) at 4 °C overnight. The membrane was washed three times with 1× TBST for 10 min, then incubated with anti-rat IgG horseradish-peroxidase (HRP)-linked antibody (Cell Signaling Technology, 7077S) in 1× TBST containing 5% milk 1 h. The membrane was washed three times with 1× TBST for 15 min. mKlotho was then detected using SuperSignal West Pico PLUS Chemiluminescent Substrate (Thermo Scientific, 34580). ImageJ software was used for quantification of the protein bands.

To pull down endogenous human Klotho (hKlotho), HK-2 cells were washed, and total protein was extracted as described above. **KB2-Biotin** or **KB2-Scramble-Biotin** was added into 300 µg of total protein and incubated at room temperature for 30 min. The following pull-down steps and western blot were described above.

Western blot performed on protein samples before or after pull-down was indicated as “Input” or “Output”. Before pull-down, 20% of total protein was prepared as “Input” samples. The other 80% of total protein was used for the following pull-down and generation of “Output” samples.

Silver stain was performed after pull-down samples separated on SDS-polyacrylamide gel per manufacturer’s protocol (Pierce Silver Stain Kit, Thermo Scientific, 24612).

#### 2.18 Imaging experiment with fixed cells

HK-2 cells were seed in 35 mm dish and grow to 70% confluency. The media was removed, and the cells were washed with 1× PBS.

To label Klotho with peptides, the cells were incubated in 1 mL of HBSS media (Gibco, 24020-117) containing 500 nM of **KB2-TAMRA** or **KB2-Scramble-TAMRA** and 25 ng/µL of WGA Alexa Fluor™ 350 Conjugate at 37 °C for 5 min. After the incubation, the cells were washed with 1× PBS followed by fixation using 400 µL of 4% paraformaldehyde (Thermo Scientific) at room temperature for 30 min. The cells were washed with 1× PBS then stained with DAPI (1 µg/mL) in the dark. The cells were then washed with 1× PBS three times.

To label Klotho with antibody, the cells were fixed using 400 µL of 4% paraformaldehyde (Thermo Scientific) at room temperature for 30 min. The cells were washed with 1× PBS then blocked with 2% BSA (w/v) in 1× PBS at room temperature for 1 h. The cells were then incubated with Klotho (Cosmo Bio, KM2076) primary antibody (1:250 dilution) in 0.1% BSA at 4 °C overnight. Primary antibody was removed then the cells were washed with 1× PBS for three times. After washing, the cells were incubated with anti-rat IgG Alexa Fluor® 647 Conjugate (Cell Signaling Technology, 4418S) as well as DAPI (1 µg/mL) in 0.1% BSA in the dark for 1 h. The cells were then washed with 1× PBS three times.

Images were taken using a DeltaVision™ Ultra microscope equipped with a scientific Complementary Metal–Oxide–Semiconductor (sCMOS) camera. A xenon arc lamp and a solid-state illumination (InsightSSI) module were used as the light source. The acquisition of images of a given culture dish was performed at constant transmission and exposure time for each channel. TAMRA was imaged at orange channel ($\lambda_{ex/em}$ = 542/597 nm) with transmission 50% and exposure time 0.2 s. WGA and DAPI were imaged at blue channel ($\lambda_{ex/em}$ = 390/435 nm) with transmission 50% and exposure time 0.1 s. Because both WGA and DAPI was imaged at blue channel, only one tracer can be chosen for each experiment to visualize membrane (WGA) or nucleus (DAPI). Antibody was imaged at red channel ($\lambda_{ex/em}$ = 632/679 nm) with transmission 20% and exposure time 0.2 s. The microscope was operated with the aid of the Acquire Ultra software. The ImageJ software (version 2.9.0) was employed for quantification of intracellular fluorescence intensity. For each measurement, the whole cell body was selected as region of interest. The integrated fluorescence from background region was subtracted from the integrated fluorescence intensity of the cell body region. To evaluate co-localization of the sensors with organelle-specific stains, Pearson’s correlation coefficients were calculated for individual cells with the JACoP plugin of ImageJ, choosing in each case the same background fluorescence signal as threshold.

#### 2.19 Imaging experiments with live cells

HK-2 cells were seed in 35 mm dish and grow to 70% confluency. The cells were treated with 500 nM of **KB2-TAMRA** and 25 ng/µL of WGA at 37 °C for 5 min in the growth media. The cells were then washed with 1× PBS three times. Images were immediately taken after washing. The following imaging steps were described above.

#### 2.20 Transfection in HK-2 cells

The KL siRNA was transfected into cells with Lipofectamine RNAiMAX reagent (Invitrogen) per the manufacturer's protocol to final concentration of 0.1 μM. Briefly, HK-2 cells were seed in 35 mm dish and grow to 50% confluency. Then, 7.5 μL of RNAiMAX reagent was diluted in 125 μL of Opti-MEM medium. Similarly, KL siRNA was diluted in 125 μL of Opti-MEM medium. Diluted siRNA was added into diluted RNAiMAX reagent and incubated at room temperature for 5 min. The total 250 μL solution was added into 35 mm dish and incubated at 37 °C for 2 days before treatment of peptides of interest or western blot (Figure S12). Cells added with solution containing RNAiMAX reagent only was considered as mock transfected.

### 3. LC-MS characterization of peptides

#### 3.1 Monomeric peptide

Name: **Pep-10**
Sequence**:** PMASDPLGVVRPRARM
HPLC: method C
HRMS (ESI-QTOF): Calcd. for $(M + H)^{1+}$: 1752.01, found: 1751.95.
**Total ion chromatogram:**

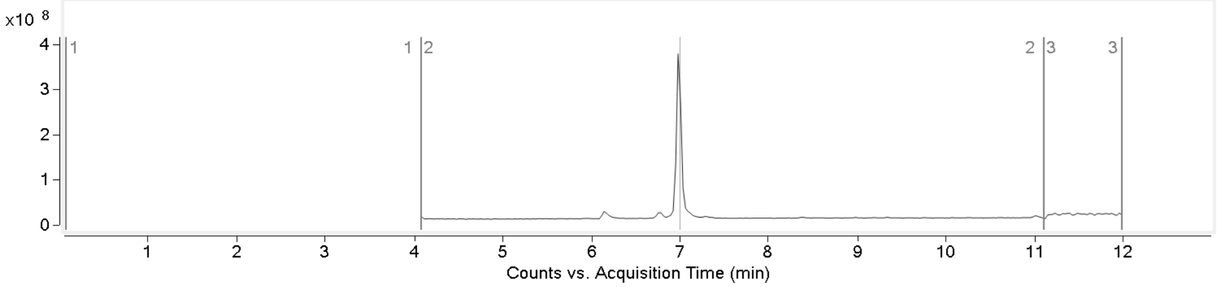


**Mass spectrum of major compound:**

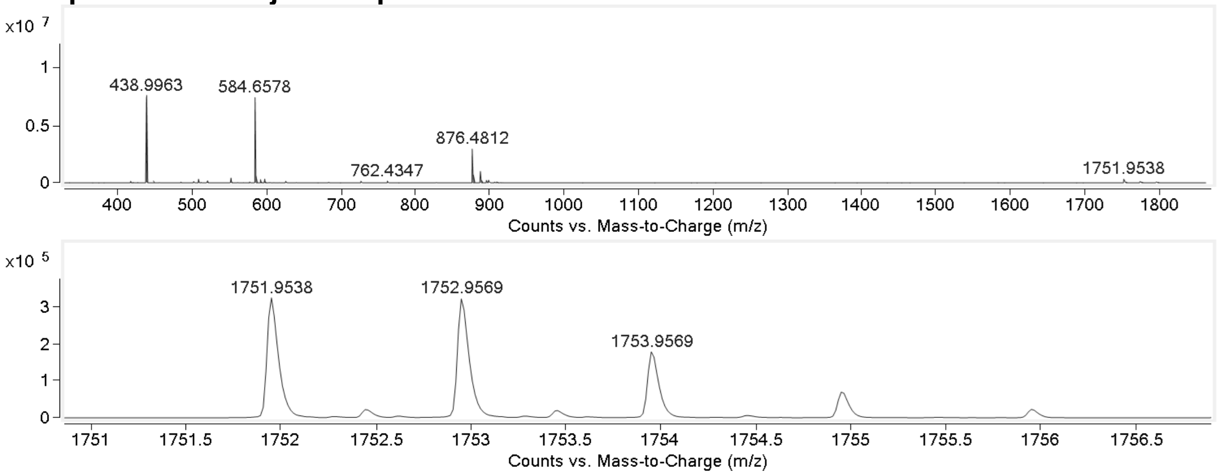


Name: **KB1-Biotin**
HPLC: method C
HRMS (ESI-QTOF): Calcd. for $(M + 2H)^{2+}$: 1083.60, found: 1083.59
**Total ion chromatogram:**

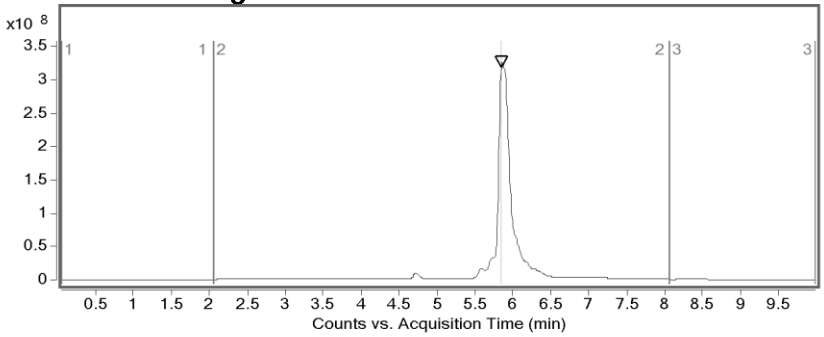


**Mass spectrum of major compound:**

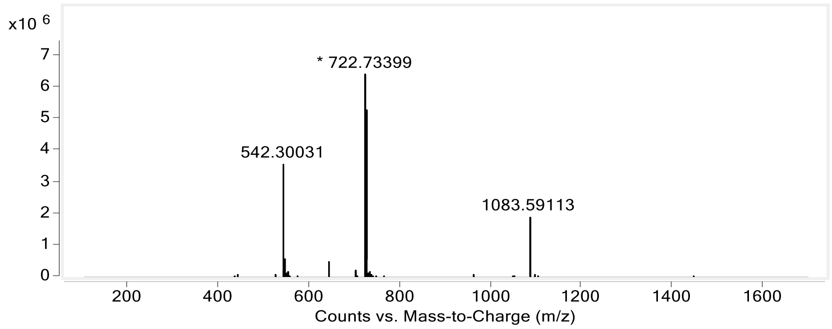


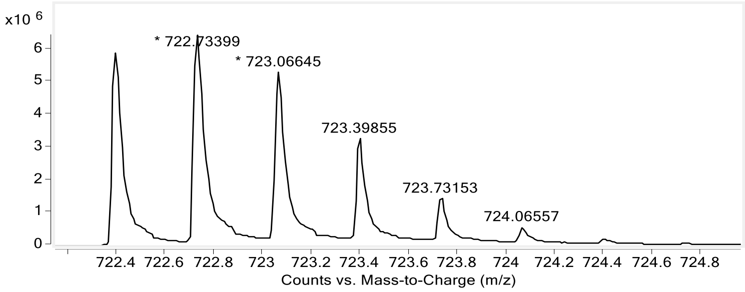


Name: **Pep10-Biotin**
Sequence: Biotin-PEG4-PMASDPLGVVRPRARM
HPLC: method D
HRMS (ESI-QTOF): Calcd. for $(M + 2H)^{2+}$:1113.11, found: 1113.09

**Total ion chromatogram:**

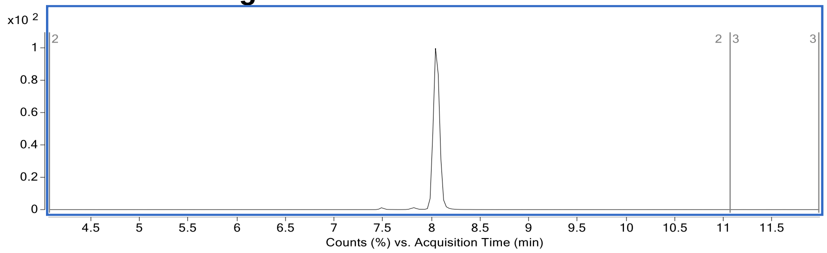


**Mass spectrum of major compound:**

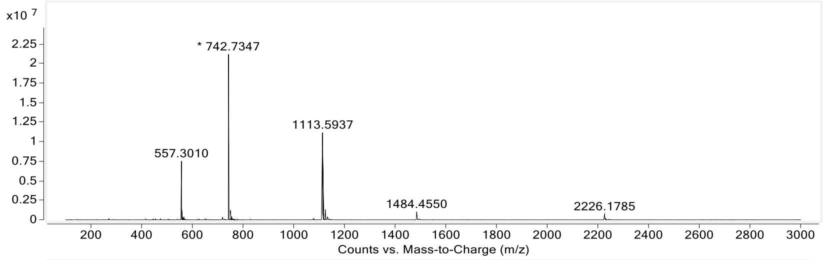


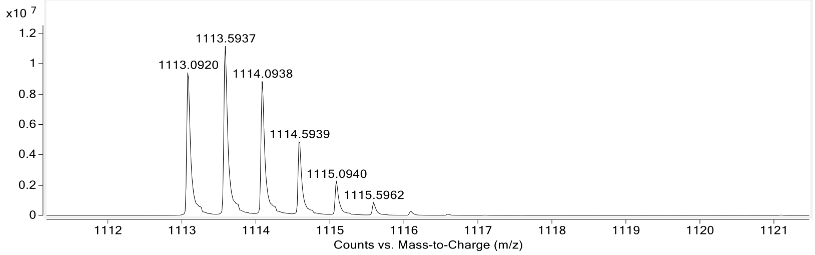

#### 3.2 Dimeric peptide

Name: **KB2**
HPLC: method A
HRMS (ESI-QTOF): Calcd. for $(M + 6H)^{6+}$: 677.76, found: 677.71.
**Total ion chromatogram:**

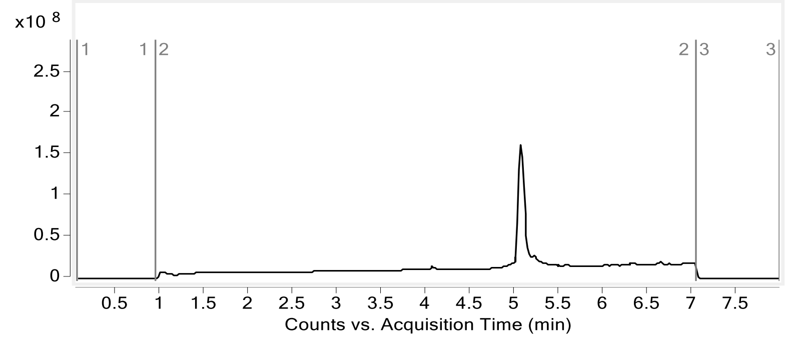


**Mass spectrum of major compound:**

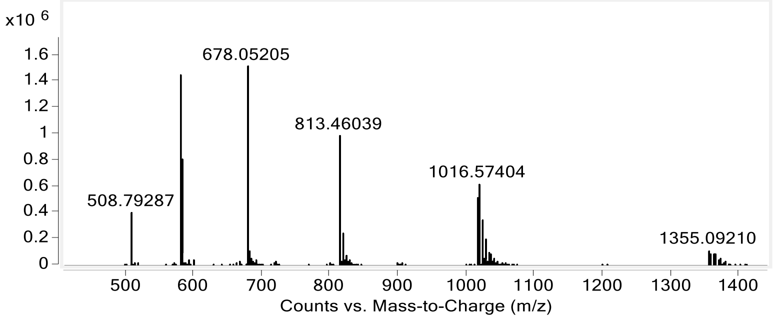


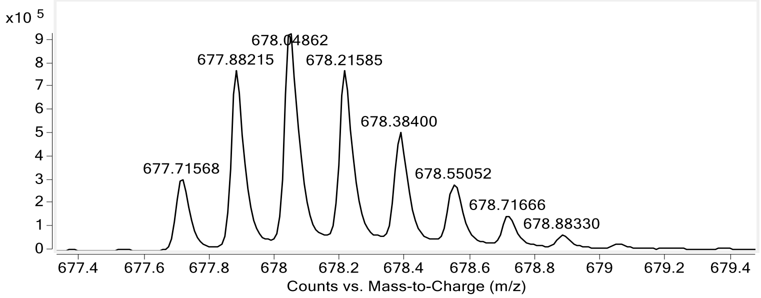


Name: **KB2-Biotin**
HPLC: method A
HRMS (ESI-QTOF): Calcd. for $(M + 4H)^{4+}$: 1134.29, found: 1134.25
**Total ion chromatogram:**

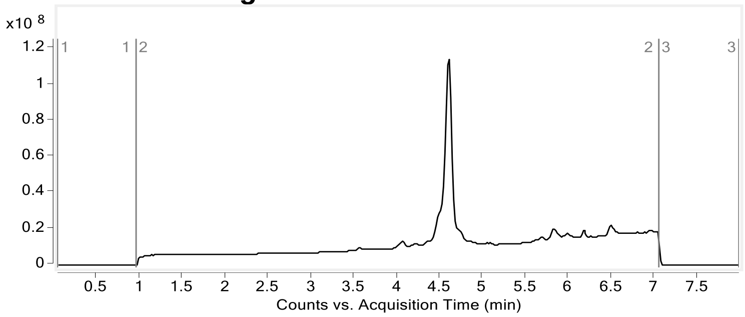


**Mass spectrum of major compound:**

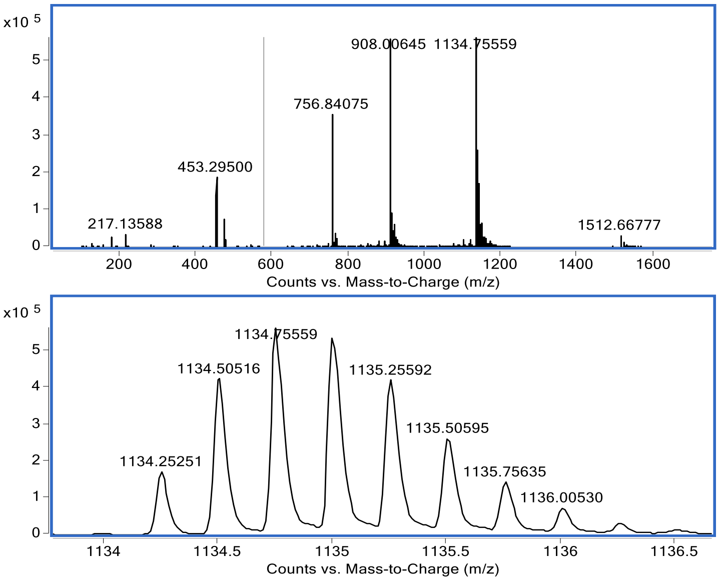


Name: **KB2-TAMRA**

HPLC: method A

HRMS (ESI-QTOF): Calcd. for $(M + 6H)^{6+}$: 746.38, found:746.40

**Total ion chromatogram:**

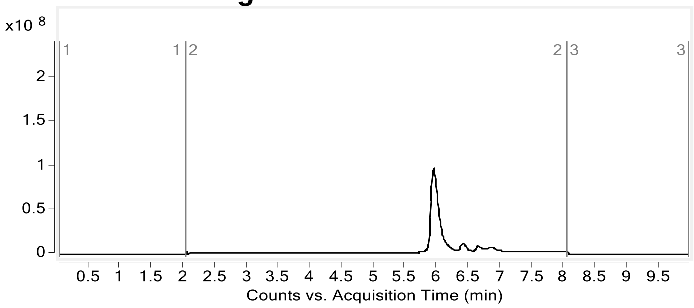


**Mass spectrum of major compound:**

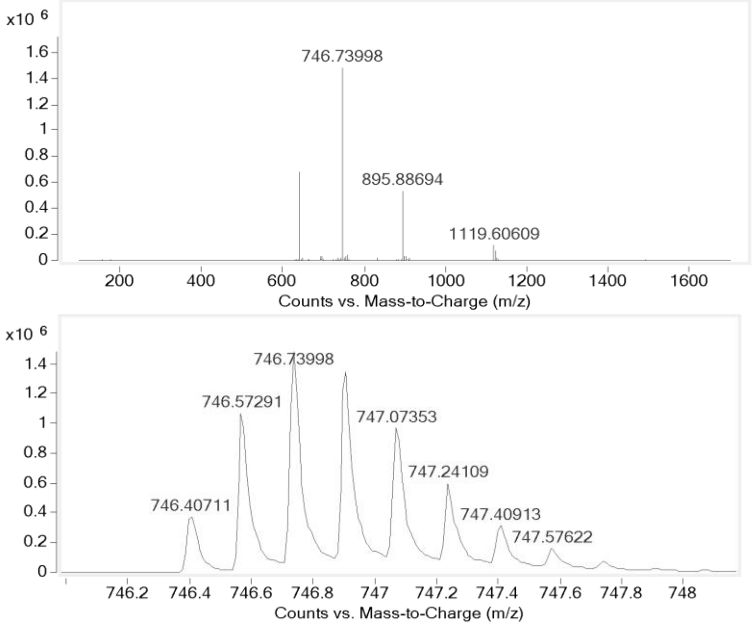

Name: **KB2-Scramble-TAMRA**
HPLC: method A
HRMS (ESI-QTOF): Calcd. for $(M + 6H)^{6+}$: 788.60, found: 788.64.
**Total ion chromatogram:**

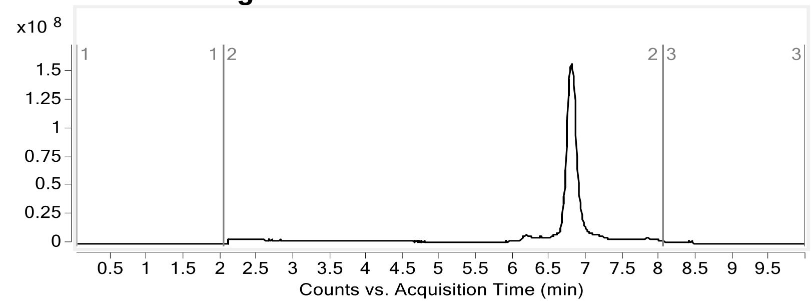


**Mass spectrum of major compound:**

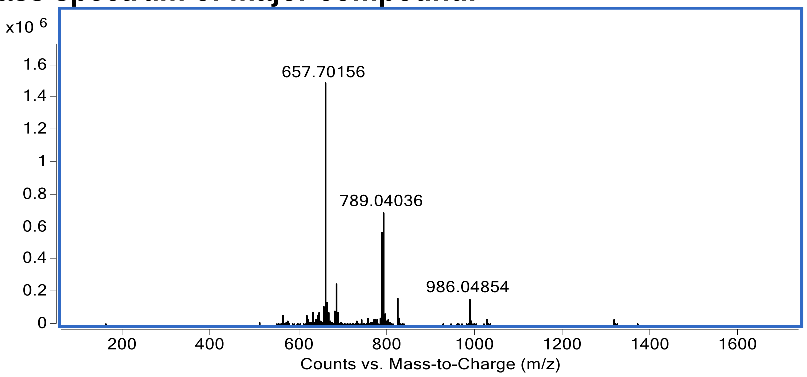


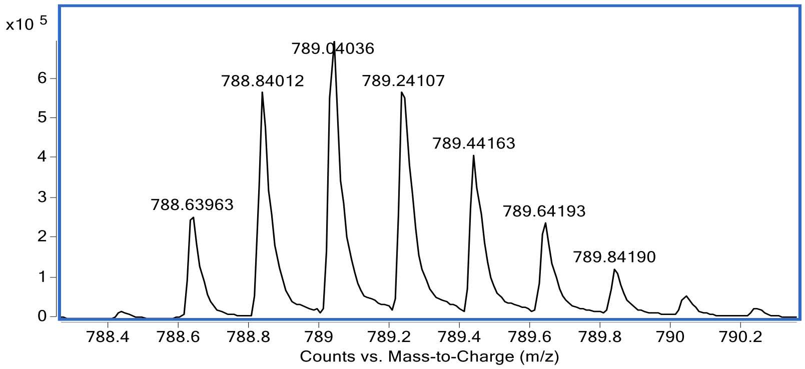


Name: **KB2-Scramble-Biotin**
HPLC: method A
HRMS (ESI-QTOF): Calcd. for $(M + 6H)^{6+}$: 726.89 , found: 726.89.
**Total ion chromatogram:**

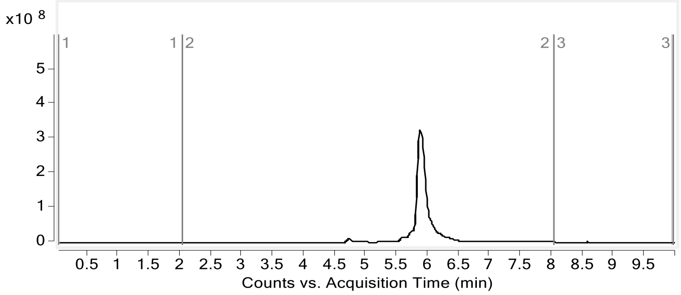


**Mass spectrum of major compound:**

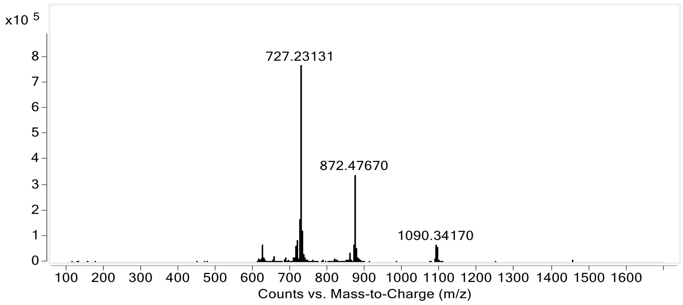


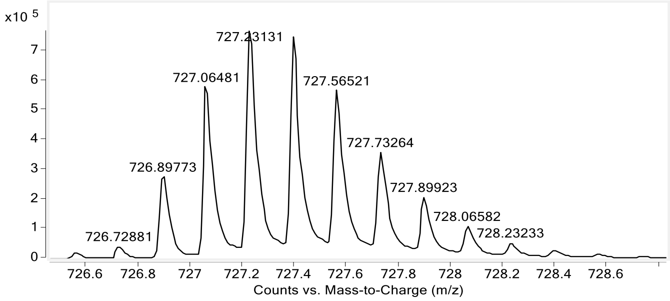


Name: **KB2-Scramble**
HPLC: method A
HRMS (ESI-QTOF): Calcd. for $(M + 5H)^{5+}$: 872.89, found: 872.47.
**Total ion chromatogram:**

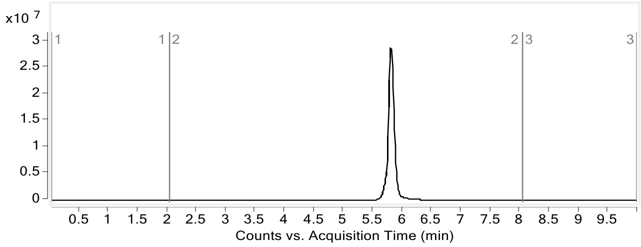


**Mass spectrum of major compound**

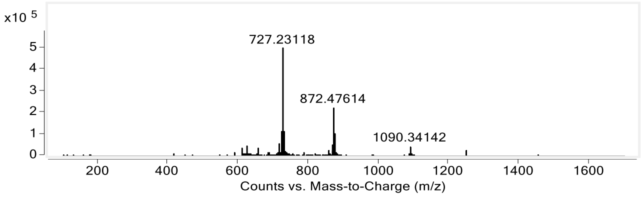


#### 3.3 Trimeric peptide

Name: **KB3**
HPLC: method A
HRMS (ESI-QTOF): Calcd. for $(M + 3H)^{3+}$: , found: 872.47.
**Total ion chromatogram:**

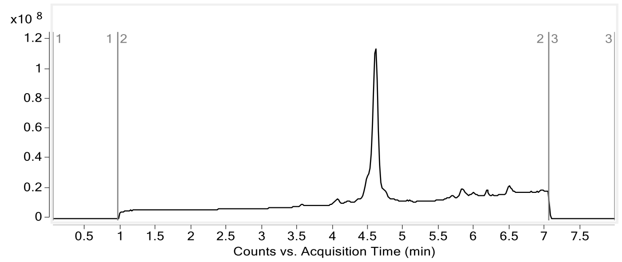


**Mass spectrum of major compound:**

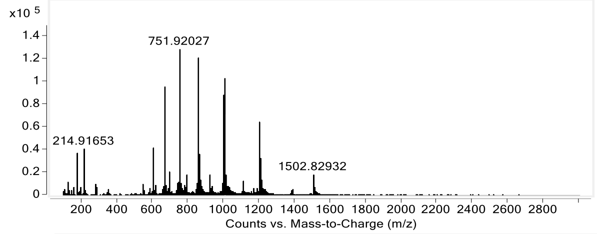


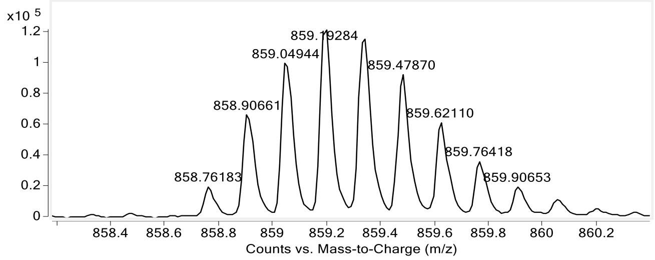


Name: **KB3-Biotin**

HPLC: method A

HRMS (ESI-QTOF): Calcd. for $(M + 6H)^{6+}$: 1087.61, found: 1087.57.

**Total ion chromatogram:**

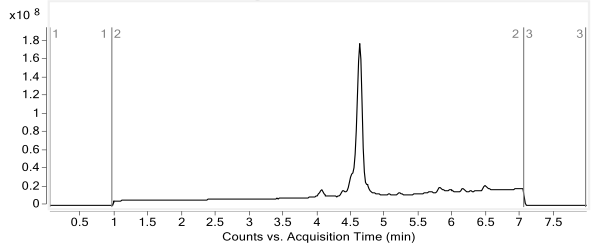


**Mass spectrum of major compound:**

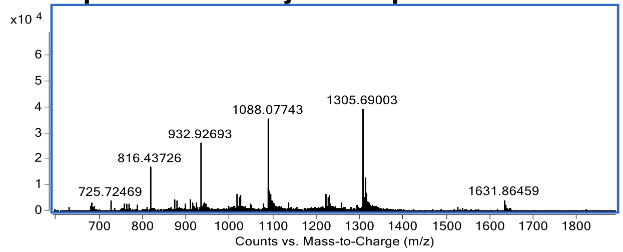

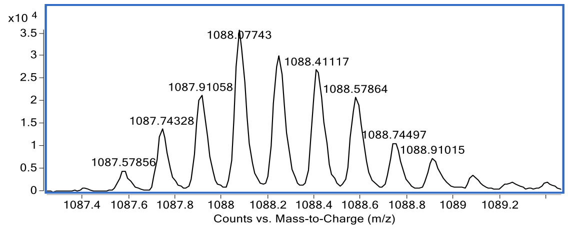


#### 3.4 Tetrameric peptide

Name: **KB4**
HPLC: method A
HRMS (ESI-QTOF): Calcd. for $(M + 6H)^{6+}$: 994.90, found: 994.64.
**Total ion chromatogram:**

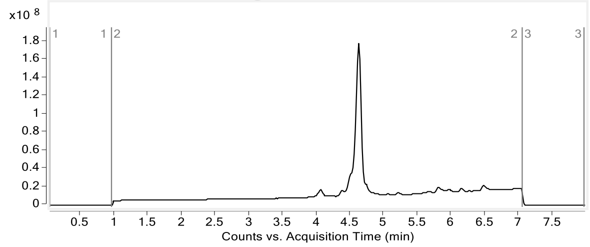


**Mass spectrum of major compound:**

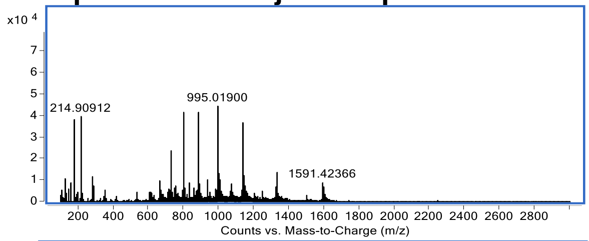


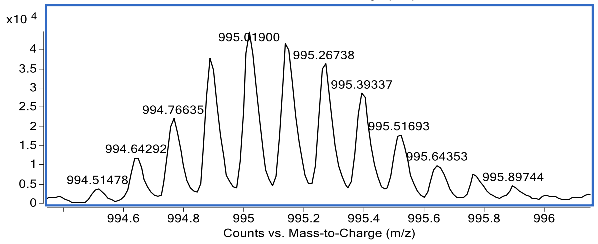


Name: **KB4-Biotin**
HPLC: method A
HRMS (ESI-QTOF): Calcd. for $(M + 6H)^{6+}$: 1346.88, found: 1346.71
**Total ion chromatogram:**

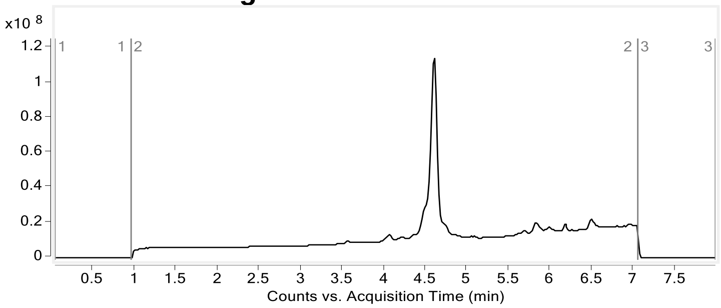

**Mass spectrum of major compound:**

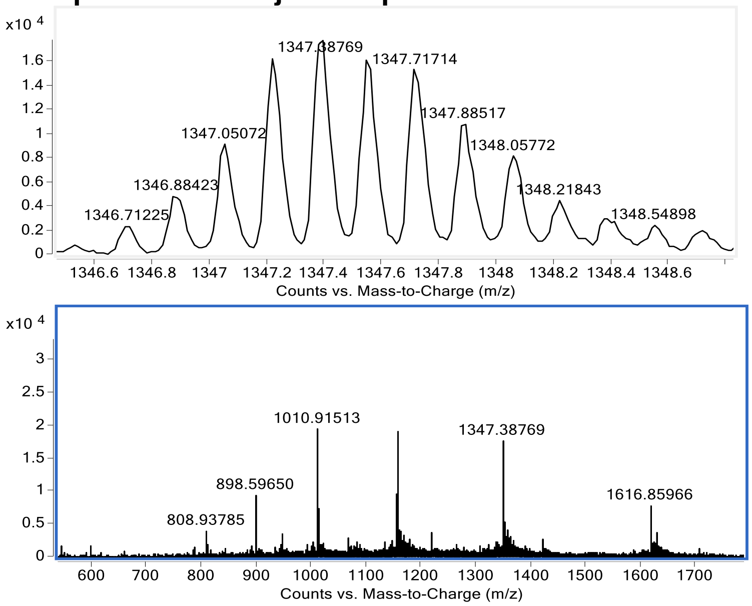
